# Targeting the TCA cycle can ameliorate widespread axonal energy deficiency in neuroinflammatory lesions

**DOI:** 10.1101/2023.04.03.535354

**Authors:** Yi-Heng Tai, Daniel Engels, Giuseppe Locatelli, Ioanna Emmanouilidis, Caroline Fecher, Delphine Theodorou, Stephan A. Müller, Simon Licht-Mayer, Mario Kreutzfeldt, Ingrid Wagner, Natalia Prudente de Mello, Sofia-Natsouko Gkotzamani, Laura Trovo, Arek Kendirli, Almir Aljovic, Michael O. Breckwoldt, Ronald Naumann, Florence M. Bareyre, Fabiana Perocchi, Don Mahad, Doron Merkler, Stefan F. Lichtenthaler, Martin Kerschensteiner, Thomas Misgeld

## Abstract

Inflammation in the central nervous system (CNS) can impair the function of neuronal mitochondria and contributes to axon degeneration in the common neuroinflammatory disease multiple sclerosis (MS). Here we combine cell type-specific mitochondrial proteomics with *in vivo* biosensor imaging to dissect how inflammation alters the molecular composition and functional capacity of neuronal mitochondria. We show that neuroinflammatory lesions in the mouse spinal cord cause widespread and persisting axonal ATP deficiency, which precedes mitochondrial oxidation and calcium overload. This axonal energy deficiency is associated with impaired electron transport chain function, but also an upstream imbalance of tricarboxylic acid (TCA) cycle enzymes, with several, including key rate-limiting, enzymes being depleted in neuronal mitochondria in experimental models and in MS lesions. Notably, viral overexpression of individual TCA enzymes can ameliorate the axonal energy deficits in neuroinflammatory lesions, suggesting that TCA cycle dysfunction in MS may be amendable to therapy.

## INTRODUCTION

Multiple sclerosis (MS) is a common neurological disease in which CNS inflammation results in myelin loss and progressive neurodegeneration. While the importance of such neurodegenerative processes for long-term disability of MS patients is well established^1–3^, the mechanistic links between CNS inflammation and neurodegeneration remain to be resolved. Emerging evidence from MS patients and models indicates that neuronal mitochondria could be critical hubs that render inflammatory signals into neurodegenerative consequences. This concept is not only supported by the essential function of mitochondria in neuronal energy homeostasis^4^, but also by previous studies showing: (i) that damaged mitochondria accumulate in axons located in experimental and human neuroinflammatory lesions^5–7^, (ii) that neuronal mitochondria acquire DNA deletions over the course of the disease^8^, and (iii) that such inflammation-induced mitochondrial damage impairs electron transport chain (ETC) function^2, 9^.

Mitochondrial damage could thus be initiated already in the highly inflamed lesions of early MS and then further amplify over the course of the disease. This process would provide a link not only between inflammation and neurodegeneration, but also between initial relapsing-remitting and later progressive pathology^2, 9^. As a result of these insights, mitochondria have emerged as a promising therapy target to prevent neurodegeneration throughout the disease course of MS. Unfortunately, so far, such strategies have failed to provide robust clinical benefits, at least in large clinical trials^10^. One of the reasons underlying this failure is that our understanding of how the molecular machinery and functional capability of mitochondria is impaired in neuroinflammatory lesions is still incomplete. Moreover, most studies so far have focused on ETC impairments^8, 11^, which might be hard to target therapeutically given the complex structure of the underlying macromolecular complexes and their genetics.

Here we apply novel *in vivo* imaging strategies combined with selective ex vivo proteomic analysis of neuronal mitochondria in a murine MS model (experimental autoimmune encephalomyelitis, EAE) to investigate the molecular underpinnings and functional consequences of inflammation-induced mitochondrial pathology. We reveal that widespread axonal ATP deficiency is initiated in acute neuroinflammatory lesions and persists in the chronic disease stage. These bioenergetic deficits precede mitochondrial redox or calcium dyshomeostasis. Selective *MitoTag*-based proteomic analysis of neuronal mitochondria isolated from acute and chronic EAE spinal cords instead revealed an imbalance of critical tricarboxylic acid (TCA) cycle enzymes, with a prominent loss of several key enzymes, including isocitrate dehydrogenase 3 (Idh3) and malate dehydrogenase 2 (Mdh2). Notably, the depletion of these key TCA cycle enzymes in neuronal mitochondria is also apparent in human MS lesions. Finally, we show that viral gene therapy, overexpressing the catalytic subunit of Idh3 or Mdh2, can partially reverse axonal ATP deficits in neuroinflammatory lesions. Our study thus provides a refined understanding of how the molecular composition of neuronal mitochondria changes in response to neuroinflammation. We identify the depletion of TCA cycle enzymes as a critical mediator of the axonal energy crisis that occurs in neuroinflammatory lesions, thus defining a potential target for therapeutic intervention.

## RESULTS

### Pervasive and persistent axonal ATP deficits emerge early in neuroinflammatory lesions

As ATP production is a central function of mitochondria that is key in axons^12^, we established an *in vivo* imaging approach to record the ATP/ADP ratio of individual axons in neuroinflammatory lesions. We generated *Thy1*-PercevalHR mice, in which neurons express PercevalHR, a genetically encoded excitation ratiometric biosensor that monitors the cytoplasmic ATP/ADP ratio^13^. Histological characterization confirmed the targeting of the sensor to a broad range of neuronal populations in the brain and spinal cord (**Extended Data Figure 1a**). To assess whether the sensor was indeed capable of recording ATP deficits in spinal axons, we pharmacologically interfered with ATP production and recorded the ATP/ADP ratio by measuring fluorescence emission after excitation at 950 and 840 nm using *in vivo* 2-photon microscopy. Application of either the glycolysis inhibitor IAA (10 mM) or the mitochondrial uncoupling agent CCCP (100 µM) to the exposed dorsal spinal cord resulted in a swift reduction of the ATP/ADP ratio in spinal axons (**Extended Data 1b,c**).

To assess the emergence of axonal energy deficits in neuroinflammatory lesions, we next induced EAE, a widely used model of MS, by immunizing *Thy1*-PercevalHR mice with myelin-oligodendrocyte glycoprotein (MOG). *In vivo* 2-photon imaging then allowed us to record the ATP/ADP ratio in axons that cross through a neuroinflammatory lesion and at the same time to stage axonal morphology, a predictor of axonal fate as we established previously^6^. We found widespread axonal ATP deficits that not only affected axons that were swollen (stage 1 axons) or fragmented (stage 2) and had hence already entered the focal axonal degeneration (FAD) process^6, 14^, but also morphologically normal axons (stage 0; **Figure 1a-e**). Axons were not only showing a reduced ATP/ADP ratio in acute lesions (analyzed 2 or 3 days after EAE symptom onset), but ATP deficits persisted in axons of all FAD stages in chronic lesions (analyzed an additional 20 days later; **Figure 1d,e**). To explore, whether neurons in their entirety lacked ATP, or whether this was a local axonal deficit, we traced individual sensory axons from the lesion back into the dorsal roots, where neuroinflammation is typically not present in EAE. Comparing the ATP/ADP ratios in the same axons close to and far from the intra-spinal lesion area revealed that ATP deficits were most pronounced at the sites of CNS inflammation (**Figure 1f-g**). To independently corroborate these results with another biosensor, we intravenously injected recombinant adeno-associated virus (rAAV.PHP.eB) vectors to neuronally express ATeam, a fluorescence resonance energy transfer (FRET)-based ATP level sensor (rAAV.hSyn:ATeam)^15^. In vivo imaging of ATeam-expressing axons confirmed a pervasive reduction in ATP availability in axons within spinal EAE lesions (**Extended Data Figure 2a-d**), while in converse, a pH biosensor (SypHer3s; rAAV.hSyn:SypHer3s)^16^, failed to identify changes in axoplasmic pH that could interfere with the PercevalHR and ATeam sensors (**Extended Data Figure 2e**). Hence, our data indicates that axons in neuroinflammatory lesions exist in a sustained state of reduced ATP availability, a finding that is in line with the localized accumulation and damage of mitochondria that we previously described in spinal cord lesions of the same MS model^6, 17^.

**Figure 1:**
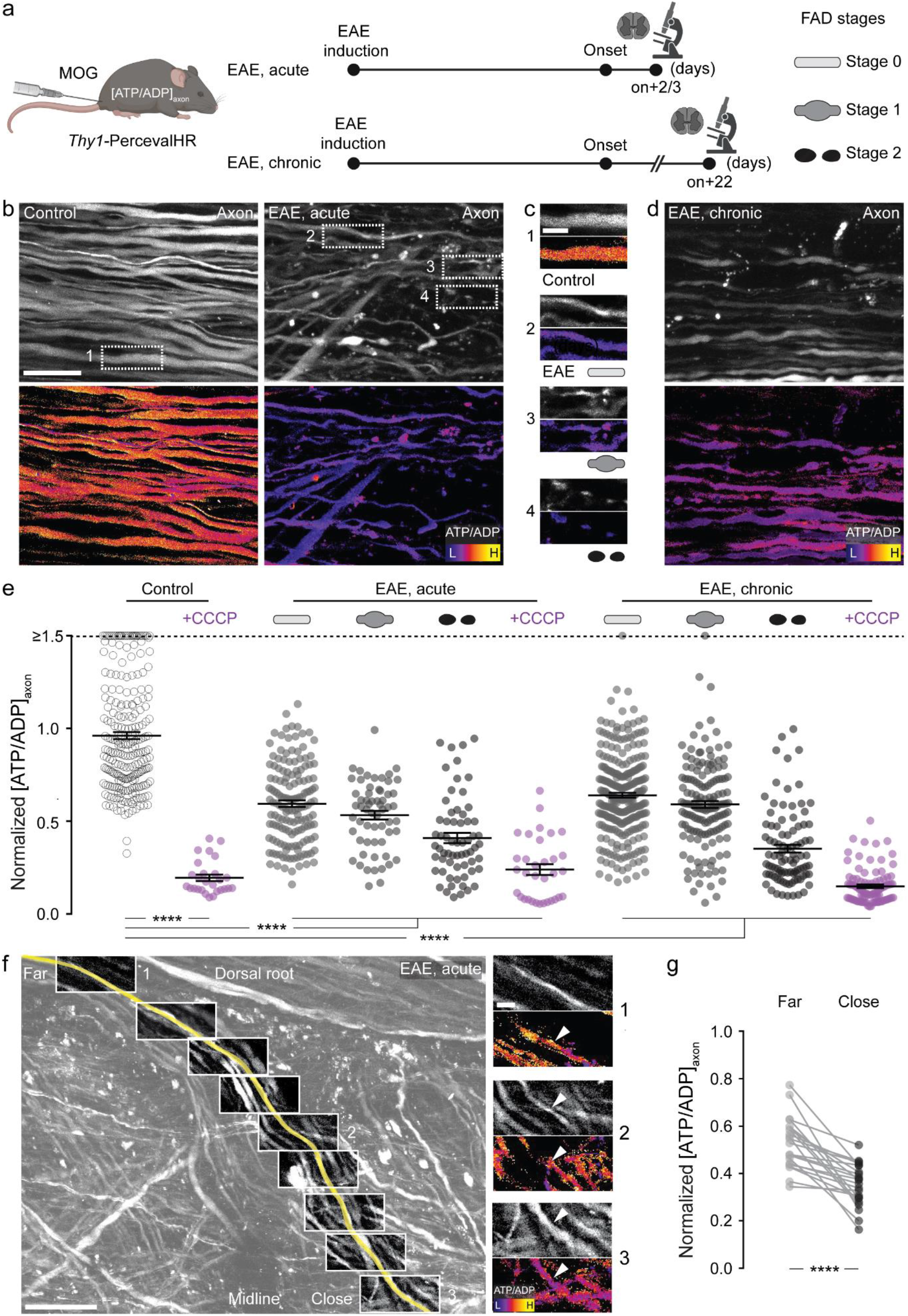
Early and pervasive axonal ATP deficits in EAE lesions. (**a**) Experimental design for axonal ATP/ADP ratio ([ATP/ADP]_axon_) measurements in experimental autoimmune encephalomyelitis (EAE); FAD: focal axonal degeneration; on: onset. (**b**) Maximum intensity projections of *in vivo* multi-photon image stacks of spinal cord axons of control (left) and acute EAE (right) in *Thy1*-PercevalHR mice. **Top**: Grayscale look-up table (LUT; λ_ex_ 950 nm). **Bottom**: Ratiometric [ATP/ADP]_axon_ LUT (λ_ex_ ratio 950 nm/840 nm). (**c**) Details from b. **Top to bottom**: [ATP/ADP]_axon_ images of control axon in healthy spinal cord, and normal-appearing, swollen, and fragmented axons in acute EAE. (**d**) Spinal cord axons during chronic EAE shown as in b. (**e**) [ATP/ADP]_axon_ of single axons in healthy and EAE mice normalized to mean of controls (magenta: axons after CCCP, 100 μM; mean ± s.e.m.; average 40 axons analyzed per mouse, in total > 800 axons, 6 control and 4 EAE, acute and 6 EAE, chronic mice; compared by Kruskal-Wallis and Dunn’s multiple comparison test; values ≥ 1.5 are lined up on the “≥1.5” dashed line). (**f**) [ATP/ADP]_axon_ gradient in a dorsal root axon traced through a lesion using *in vivo* multiphoton imaging. **Left**: Traced axon pseudo-colored yellow, running from root at top left to the lesion center at lower right – superimposed on full volume projection, grayscale LUT, λ_ex_ 950 nm. Boxes: Locations of high-resolution stacks. **Right**: Details showing the [ATP/ADP]_axon_ gradient between locations far from (1) vs. close to the lesion (3). LUTs as in b. (**g**) Paired analysis of [ATP/ADP]_axon_ far vs. close to the lesion (n = 22 axons from 3 EAE mice; paired t-test; normalized to control). Scale bars: 25 μm in b, also applied to d; 10 μm in c and f (right); 100 μm in f (left). ***, p < 0.005; ****, p < 0.001.

### Axonal ATP deficits precede mitochondrial redox and calcium dyshomeostasis

Oxidative stress to axons had previously been proposed as a mediator of mitochondrial damage in neuroinflammatory lesions^1–3, 9, 18^. Thus, we set out to determine if and when an altered redox status of axonal mitochondria would emerge in relation to the onset of neuroinflammation-induced ATP deficits. We used *Thy1*-mitoGrx-roGFP mice, which allow recording the glutathione redox potential in neuronal mitochondria, as we previously established^19^. After crossing these redox reporter mice to *Thy1-*OFP mice^20^ to allow staging of axonal morphology, we induced EAE and performed ratiometric *in vivo* confocal imaging using single-photon excitation at 405 and 488 nm at the peak of acute disease symptoms. An oxidative shift in mitochondrial redox state was only apparent in fragmented (stage 2) axons, while mitochondria in normal-appearing (stage 0) and swollen (stage 1) axons were unaltered compared to control axons in a healthy spinal cord (**Figure 2a-d**). This is notable, as the analysis of individual mitochondria confirmed that damage-related mitochondria shape changes, such as rounding up (reduced shape factor), are already present at the onset of FAD (**Figure 2e**)^6, 17^. Decompensated mitochondrial redox stress thus seems to be a late event during inflammatory axon degeneration and affects only a small proportion of non-fragmented axons in neuroinflammatory lesions. Oxidative mitochondrial damage per se is hence unlikely to underlie the widespread axonal ATP deficits in EAE.

**Figure 2:**
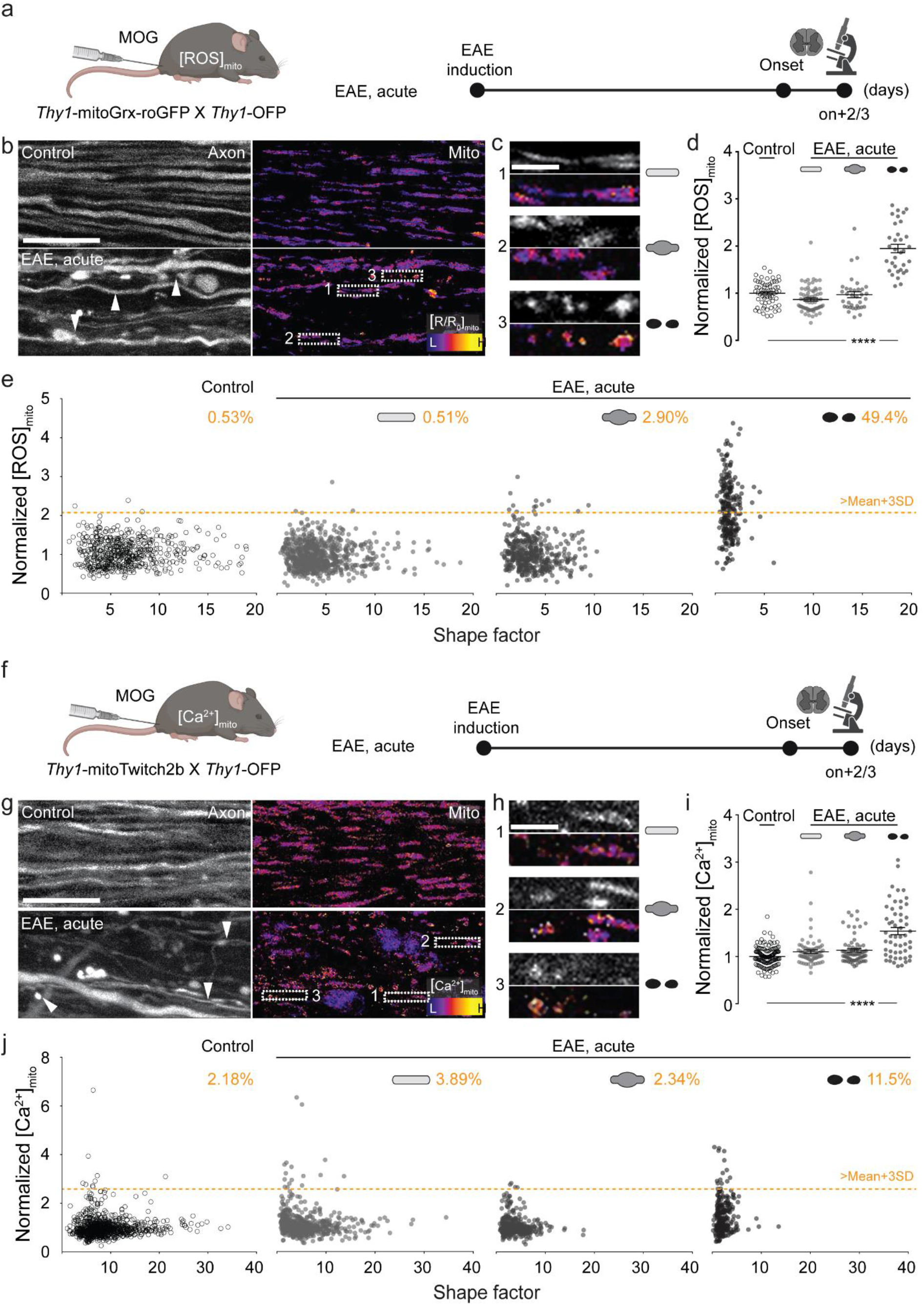
Late changes in axonal ROS and calcium homeostasis in EAE lesions. (**a**) Experimental design to measure mitochondrial ROS ([ROS]_mito_) in EAE. (**b**) Maximum intensity projections of *in vivo* confocal image stacks of spinal cord axons of control (top) and acute EAE (bottom) in *Thy1*-mitoGrx-roGFP ⊆ *Thy1*-OFP mice. **Left**: OFP channel shown with grayscale LUT. **Right**: Ratiometric LUT of [ROS]_mito_ (λ_ex_ ratio 405 nm/488 nm). (**c**) Details from b. **Top to bottom**: mitochondria morphologies and [ROS]_mito_ in normal-appearing, swollen and fragmented axons in acute EAE; grayscale LUT of OFP channel above ratiometric [ROS]_mito_ images. (**d**) Average [ROS]_mito_ of single axons in control and acute EAE mice normalized to mean of controls (mean ± s.e.m.; average 17 axons analyzed per mouse, in total > 200 axons, 5 control and 6 EAE mice compared by one-way ANOVA and Tukey’s multiple comparison test). (**e**) Single organelle correlation analysis of mitochondrial shape factor (length/width ratio) and [ROS]_mito_ of axons plotted in d. Percentages indicate fraction of mitochondria with [ROS]_mito_ > control mean + 3SD (orange line) in each axon stage. (**f**) Experimental design to measure mitochondrial Ca^2+^ level ([Ca^2+^]_mito_) in EAE. (**g**) Maximum intensity projections of *in vivo* multiphoton images of spinal cord axons of control (top) and acute EAE (bottom) in *Thy1*-mitoTwitch2b ⊆ *Thy1*-OFP mice. **Left**: OFP channel shown with grayscale look-up table. **Right**: Ratiometric LUT of [Ca^2+^]_mito_ (YFP/CFP emission ratio). (**h**) Details from g. **Top to bottom**: mitochondria morphologies and [Ca^2+^]_mito_ represented as in c; grayscale LUT of OFP channel above ratiometric [Ca^2+^]_mito_ images. (**i**) Average [Ca^2+^]_mito_ of single axons in control and acute EAE mice normalized to mean of controls (mean ± s.e.m.; average 16 axons analyzed, in total > 300 axons, 9 control and 11 EAE mice compared by one-way ANOVA and Tukey’s multiple comparison test). (**j**) Single organelle correlation analysis of mitochondrial shape factor (length/width ratio) and [Ca^2+^]_mito_ of axons plotted in i. Percentages indicate fraction of mitochondria with [Ca^2+^]_mito_ > control mean + 3SD (orange line) in each axon stage. Arrow heads indicate axons with different FAD stages. Scale bars: 25 μm in b and g; 10 μm in c and h. ****, p < 0.001.

In addition to oxidative damage, calcium overload was proposed to mediate mitochondrial dysfunction^1, 21^. To monitor calcium handling of axonal mitochondria *in vivo*, we generated a transgenic reporter mouse line that expresses the ratiometric calcium sensor Twitch2b^22^ selectively in neuronal mitochondria. Histological characterization of these *Thy1*-mitoTwitch2b mice confirmed widespread sensor expression in CNS neurons. Moreover, induction of localized laser lesions in the spinal cord demonstrated that the sensor can record relevant pathological calcium signals *in vivo* (**Extended Data Figure 1d-f**). We then crossed these reporter mice to *Thy1*-OFP mice to relate axonal morphology to mitochondrial calcium levels and shape in acute EAE lesions. Like our analysis of mitochondrial redox state, we found that mitochondrial calcium levels are primarily increased in fragmented (stage 2) axons, i.e., during the end stage of FAD, while normal-appearing (stage 0) and swollen (stage 1) axons showed no pronounced calcium dyshomeostasis, despite clear morphological signs of organelle damage (**Figure 2f-j**).

Taken together, these experiments indicate that overt dysregulation of the mitochondrial redox state and calcium handling are rather late events during inflammatory axon degeneration. Hence, a molecularly distinct dysregulation of mitochondrial function and molecular composition must explain the prodromal and pervasive ATP deficits in EAE axons.

### Proteomics of neuronal mitochondria reveals ETC and TCA cycle enzyme depletion

To obtain an unbiased characterization of the neuroinflammatory changes to the molecular make-up of neuronal mitochondria, we adapted a cell type-specific proteomics approach that we recently established^23^. For this, we performed intrathecal injections of a rAAV.hSyn: Cre virus into the ventricles of *MitoTag* neonates, which resulted in the tagging of the outer membrane of neuronal mitochondria with GFP. This enabled selective isolation and subsequent mass spectrometry analysis of these mitochondria from the spinal cords of healthy mice, as well as from different stages of EAE (**Figure 3a,b**, **Extended Data Figure 3a-c** and **Extended Data Table 1**), which revealed pronounced changes to the mitochondrial proteome of neurons. These changes could not be predicted by protein lifetime, location inside the mitochondria or whether a protein was encoded in the nucleus vs. the mitochondrial DNA (**Extended Data Figure 3d-f**). Gene set analysis of the major dysregulated pathways converged on the ETC and the TCA cycle, which are abundant in healthy neuronal mitochondria, but depleted in acute neuroinflammatory lesions (**Figure 3c,d** and **Extended Data Figure 3g,h**). The ETC complexes were uniformly affected, typically more strongly in acute than in chronic EAE lesions (**Figure 3d** and **Extended Data Figure 3g,h and 4a**). This corresponded with reduced axon complex IV (COX) activity as measured in situ by a histochemical assay^24^ (**Extended Data Figure 4b,c**) and corroborated the previously described neuronal ETC dysfunction in neuroinflammatory lesions^5^. Notably, however, also TCA cycle enzymes showed pronounced dysregulation (**Figure 3d,e**). To corroborate these proteomic findings in EAE, expand them to MS and probe their bioenergetic significance, we focused our further analysis on Idh3, which mediates the irreversible oxidative decarboxylation of isocitrate, and Mdh2, which oxidizes malate to oxaloacetate. Both enzymes generate NADH, which is subsequently used for ATP generation by the ETC, with Idh3 being rate-limiting for the TCA cycle, while Mdh2 catalyses a key reaction that links the TCA cycle to anaplerotic reactions^25, 26^. At the same time, Idh2, which accelerates the reversible NADP-dependent conversion of isocitrate to α-ketoglutarate, appears to be differentially affected from Idh3 and Mdh2, as its abundance is unchanged in proteomes from acute and increased in chronic neuroinflammatory lesions (**Figure 3d,e**). Expression changes of key TCA cycle related enzymes, including Idh3, Idh2 and Mdh2, were also present on the transcriptional level as revealed by analysis of published RiboTag translatomes of spinal motoneurons in EAE^27^ (**Figure 3f**). Hence, neuroinflammation caused a marked alteration in the expression of TCA cycle enzymes, including the rate-limiting enzyme Idh3, which likely hinders neuronal mitochondria from generating sufficient amounts of ATP.

**Figure 3:**
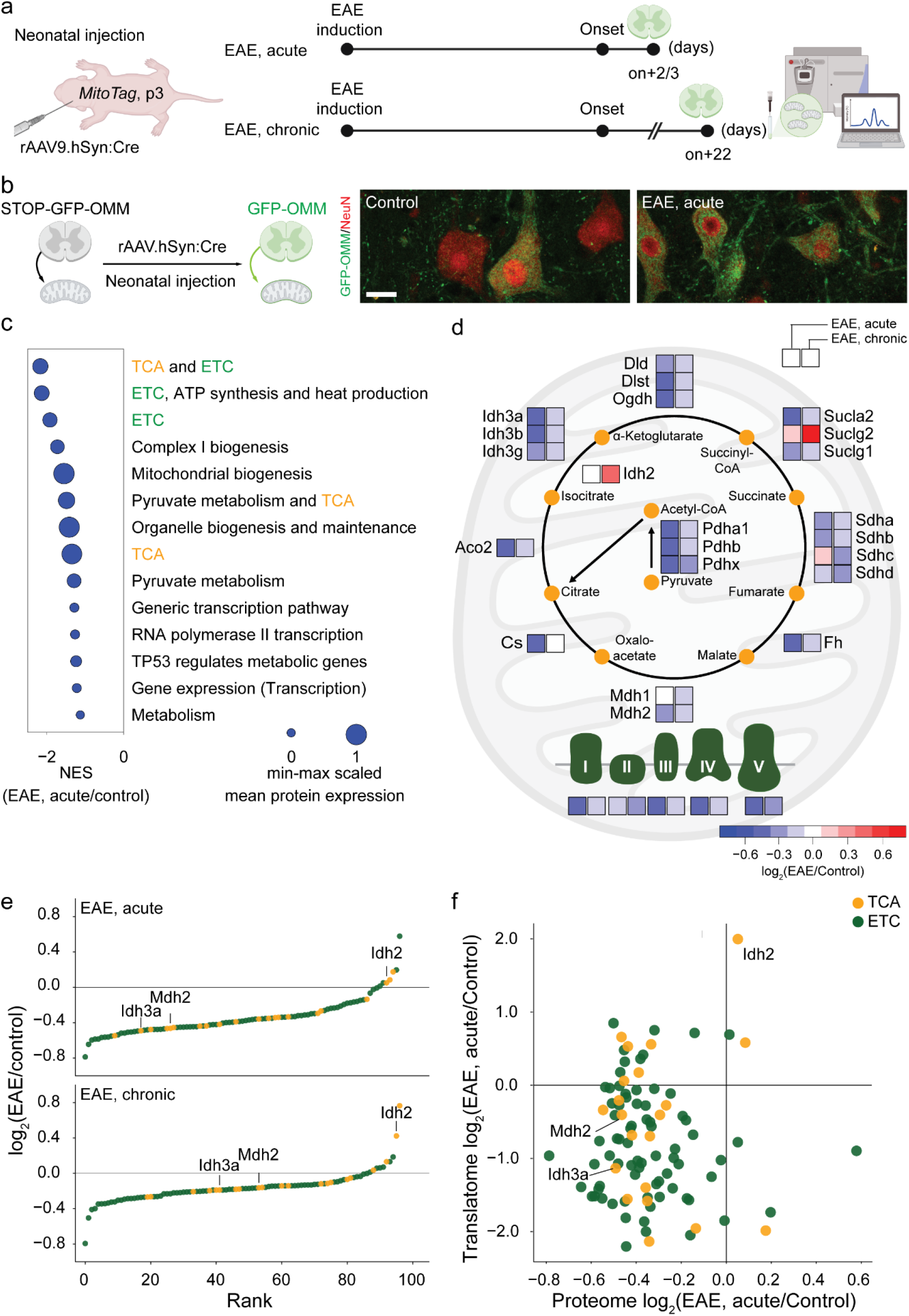
ETC depletion and TCA cycle imbalance in neurons during EAE. (**a**) Experimental design for combined AAV/*MitoTag*-based proteomic analysis of neuronal mitochondria in EAE. (**b**) Confocal image of GFP expression (green) in rAAV9.hSyn:Cre transduced neurons (NeuN, red) in a control and EAE *MitoTag* mouse spinal cord. (**c**) Annotations of the most down-regulated pathways (Reactome^59, 60^, version 7.4) in *MitoTag* proteomes of neuronal mitochondria in acute EAE. Dot size indicates min-max scaled mean protein expression level ranged from 0 to 1. (**d**) Relative abundance of the TCA cycle and ETC components in neuronal mitochondria. Average shown as color-coded log_2_(EAE/Control) for acute and chronic EAE compared to respective controls. (**e**) Rank of TCA cycle (orange) and ETC (green) proteins according to log_2_FC expression in acute (top) and chronic (bottom) EAE. (**f**) Correlation between neuronal transcript and protein levels of TCA cycle and ETC proteins in EAE. Transcriptomic data was re-analyzed from Schattling et al.^27^ Scale bars: 25 μm in b.

### Axonal Idh3 and Mdh2 are depleted not only in EAE but also in MS lesions

We next aimed to confirm the proteomics results by immunofluorescence analysis of neuronal mitochondria in EAE, as well as in MS lesions. First, we quantified the expression levels of a subset of TCA cycle enzymes, for which the neuronal *MitoTag* analysis predicted dysregulation in EAE. We performed immunostainings on spinal cord sections from *Thy1*-mitoRFP mice^19^, where analysis can be restricted to red fluorescent protein (RFP)-tagged neuronal somata mitochondria. Indeed, the levels of Idh3a (the catalytic subunit of Idh3; **Figure 4b,d**), as well as Mdh2 (**Figure 4e,g**) were decreased by more than 50 % in EAE lesion-adjacent areas compared to controls. As predicted by our proteomics analysis, the expression of Idh2 was slightly increased (**Figure 4h,j**). We further confirmed the depletion of Idh3a and Mdh2 in axons per se, the neuronal compartment where we detected ATP depletion (**Figure 4c,d,f,g**). Second, to explore whether similar expression changes of TCA cycle enzymes were also present in MS lesions, we performed immunofluorescence analysis of axonal Idh3a, Idh2 and Mdh2 in brain biopsy and autopsy sections from 7 MS patients (**Extended Data Table 2**). As fixation protocols are typically variable in patient-derived material, we internally normalized the expression of a given TCA cycle enzyme between the lesion area and the adjacent normal-appearing white matter (NAWM) on the same section (**Figure 5a**). This analysis revealed a reduction of Idh3a and Mdh2 in MS lesions (**Figure 5b-e**), which was also apparent in chronic active MS lesions, indicating that depletion of TCA cycle enzymes persists long-term. Notably, Idh2 expression was not markedly affected, as predicted from the animal model (**Figure 5f,g**). Taken together, our findings show that TCA cycle disruption is a persistent neuronal change during MS lesion formation and progression.

**Figure 4:**
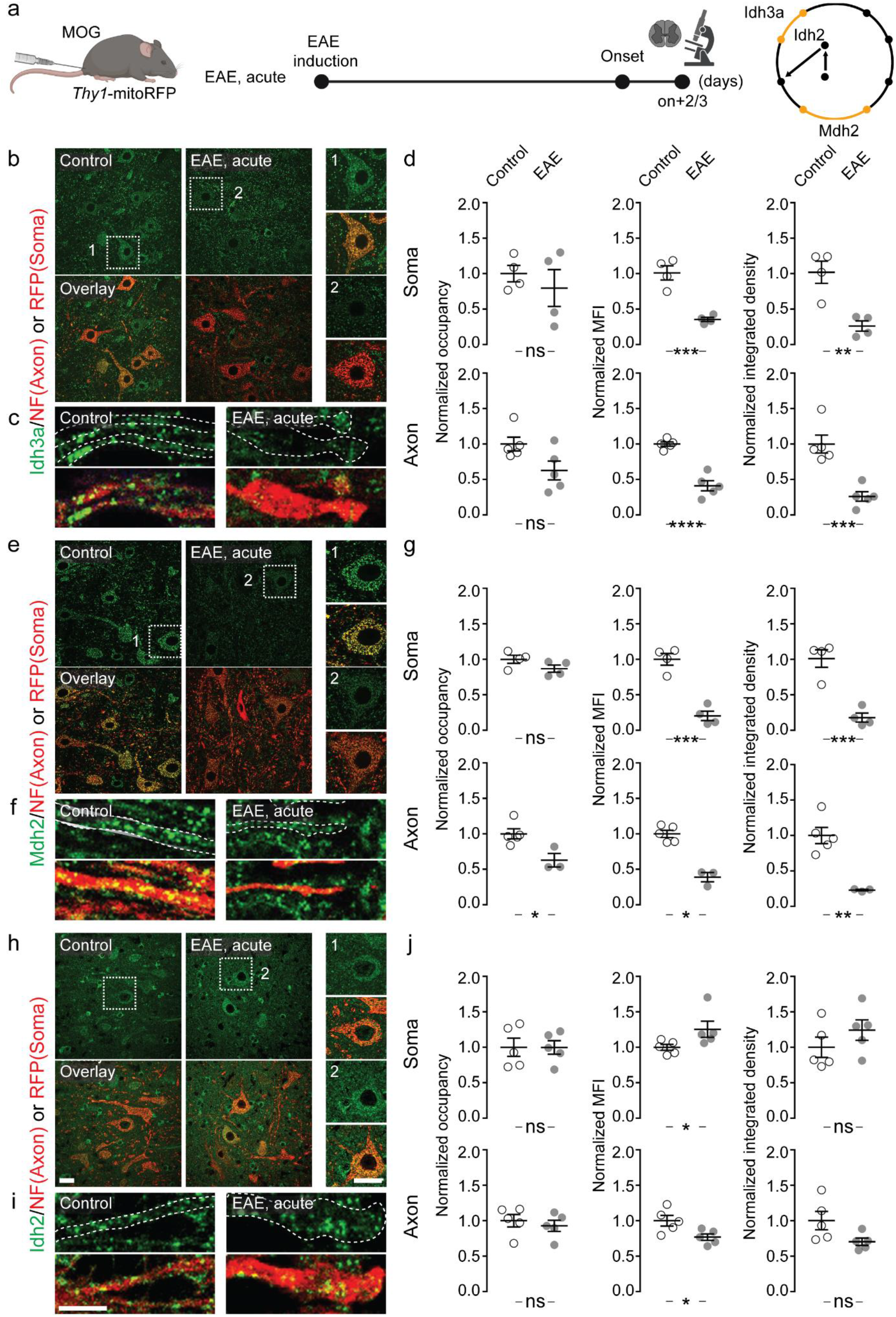
TCA cycle enzymes – Idh3, 2 and Mdh2 – are dysregulated in neurons in EAE. (**a**) Experimental design to validate dysregulated TCA cycle enzymes identified by *MitoTag* proteomics in EAE. (**b-j**) Immunofluorescence analysis of Idh3a (**b-d**), Mdh2 (**e-g**) and Idh2 (**h-j**) in spinal cord neuronal somata (top) and axons (bottom) of control (left panels) or of acute EAE (right panels) using *Thy1*-mitoRFP mice (TCA cycle enzymes, green; RFP, red for neuronal mitochondria in somata or, NF for axonal staining, respectively). Graphs (**d, g, j**) show area occupancy, mean fluorescence intensities and their product – integrated density – in EAE neuronal somata and axons normalized to the mean of control (mean ± s.e.m.; n = 4-5 mice, compared per animal by two-tailed, unpaired Student’s t-test. Scale bar: 25 μm in h, applies also to b, e, and their details; 10 μm in i, applies also to c and f. *, p < 0.05; **, p < 0.01, ***, p < 0.005; ****, p < 0.001.

**Figure 5:**
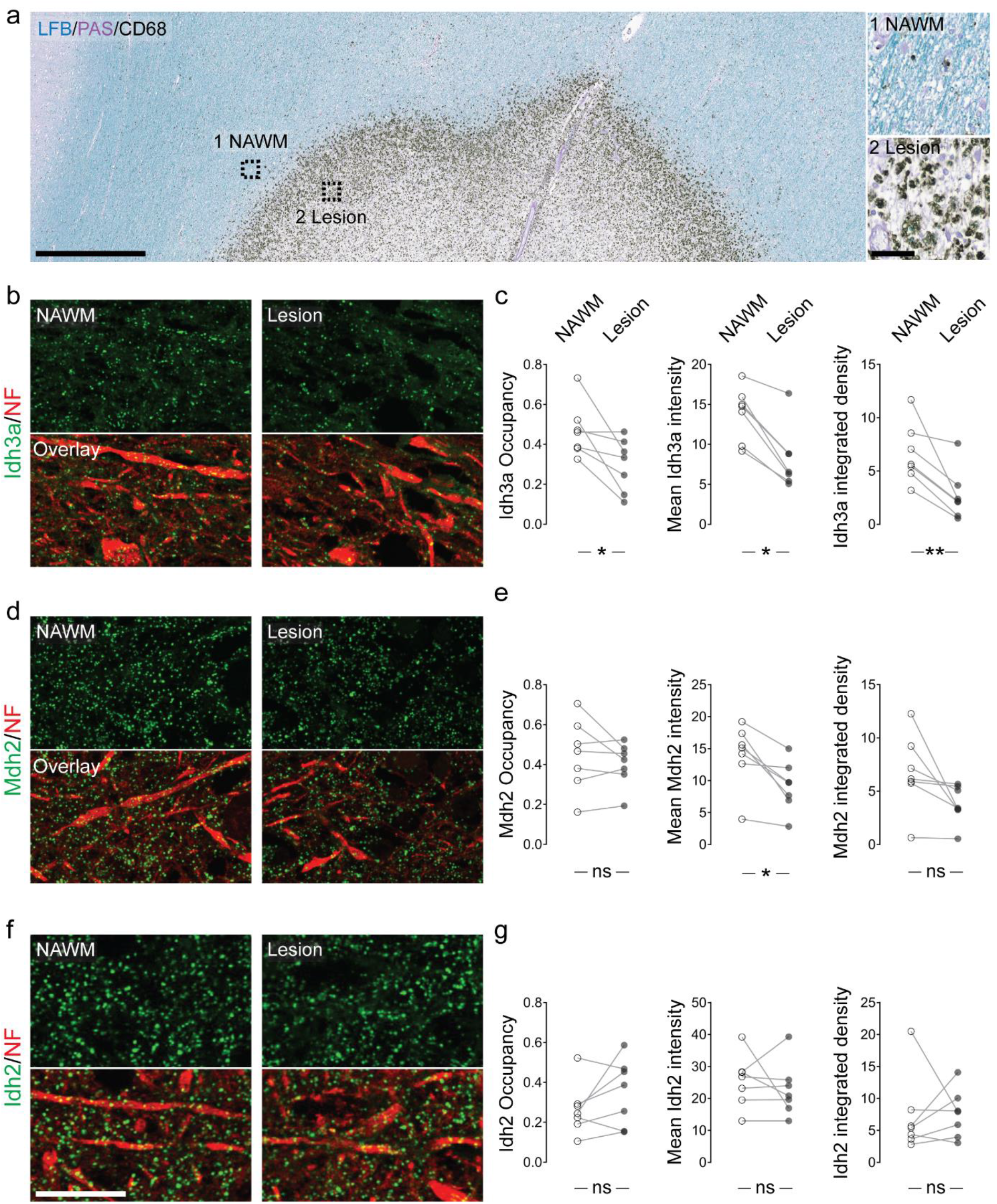
TCA cycle enzymes – Idh3, 2 and Mdh2 – are dysregulated in neurons in MS. (**a**) Overview of an MS lesion in human cortex, with areas of normal-appearing white matter (NAWM) and the lesion site marked and magnified in insets. Myelin (Luxol fast blue, LFB, blue; periodic acid–Schiff, PAS, purple) and macrophages (CD68, black) are labelled. (**b, c**) Immunofluorescence analysis of Idh3a in cortical axons in NAWM (left panels) or lesion areas (right; Idh3a, green; neurofilament, NF, red). (d, e) Immunofluorescence analysis of Mdh2 in cortical axons in NAWM (left panels) or lesion areas (right; Mdh2, green; neurofilament, NF, red). (f, g) Immunofluorescence analysis of Idh2 in cortical axons in NAWM (left panels) or lesion areas (right; Idh2, green; neurofilament, NF, red). Graphs (**c, e, g**) show area occupancy, mean fluorescence intensities and their product – integrated density – in pairs of NAWM and lesion areas from n = 7 cases, as listed in Extended Data Table 2 and compared by paired t-test. Scale bars: 1000 μm in a; 50 μm in inset; 25 μm in f, also applies to b and d. *, p < 0.05; **, p<0.01.

### Restoring Idh3a and Mdh2 expression partially rectifies axonal ATP deficits in neuroinflammatory lesions

Considering the pronounced depletion of Idh3a in EAE and MS, as well as its critical role in the TCA cycle, we explored whether overexpression of individual depleted TCA cycle components, such as Idh3a, the enzyme’s catalytic subunit^28^, could restore axonal ATP levels in neuroinflammatory lesions. We initially focused on Idh3 because of its role as a pace-maker enzymes of the TCA cycle^29^ and the substantial translational efforts that are underway to target Idh enzymes as these are mutated in several cancers^30–35^. For overexpressing Idh3a together with a fluorescent marker (tdTomato), we used the systemic injection of a rAAV.PHP.eB virus, which drives transgene expression via a pan-neuronal promoter (human Synapsin)^36^. Confocal analysis confirmed that a substantial fraction of CNS neurons was transduced with the rAAV.hSyn:Idh3a-tdTomato virus as indicated by tdTomato expression. Moreover, tdTomato-labelled neurons in EAE spinal cords showed a marked increase in mitochondrial Idh3a expression (**Extended Data Figure 5**). We then directly assessed the effects of Idh3a restoration on axonal energy deficits in neuroinflammatory lesions by inducing EAE in *Thy1*-PercevalHR mice injected with either the rAAV.hSyn:Idh3a-tdTomato virus or a control rAAV.hSyn:Cre-tdTomato virus, carrying a similarly sized, but in our setting inert transgene*. In vivo* imaging of the surgically exposed spinal cord in EAE-induced animals again confirmed strong tdTomato-expression in a fraction of axons, while other PercevalHR-expressing axons were tdTomato-negative and provided an internal control (**Figure 6a,b**). Compared to this control axon population (which rules out general effects on lesion activity by the virus), transduction with a rAAV.Syn:Idh3a-tdTomato virus significantly increased the ATP/ADP ratio and thus partially restored the axonal energy deficiency in neuroinflammatory lesions (**Figure 6c**). No such rescue was observed in axons transduced with the control rAAV.hSyn:Cre-tdTomato (**Figure 6d**). Notably, ATP/ADP ratios rose in axons of all damage stages, suggesting that correcting Idh3a levels could also counteract ATP deficiencies in more advanced stages of FAD. By extending the observation period from the acute phase after the EAE peak by three weeks, we confirmed that both the reduction in ATP/ADP ratio and a reversing effect of Idh3a overexpression persisted, albeit the latter’s effects appeared to abate **Extended Data Figure 6a-c**). Next, we asked whether the effect of overexpressing Idh3a would be unique to this enzyme by targeting Mdh2 in a similar way. We observed that viral gene transfer of Mdh2 partially restored axonal ATP/ADP levels in neuroinflammatory lesions at least in advanced stages of axon damage (**Extended Data Figure 7**).

**Figure 6:**
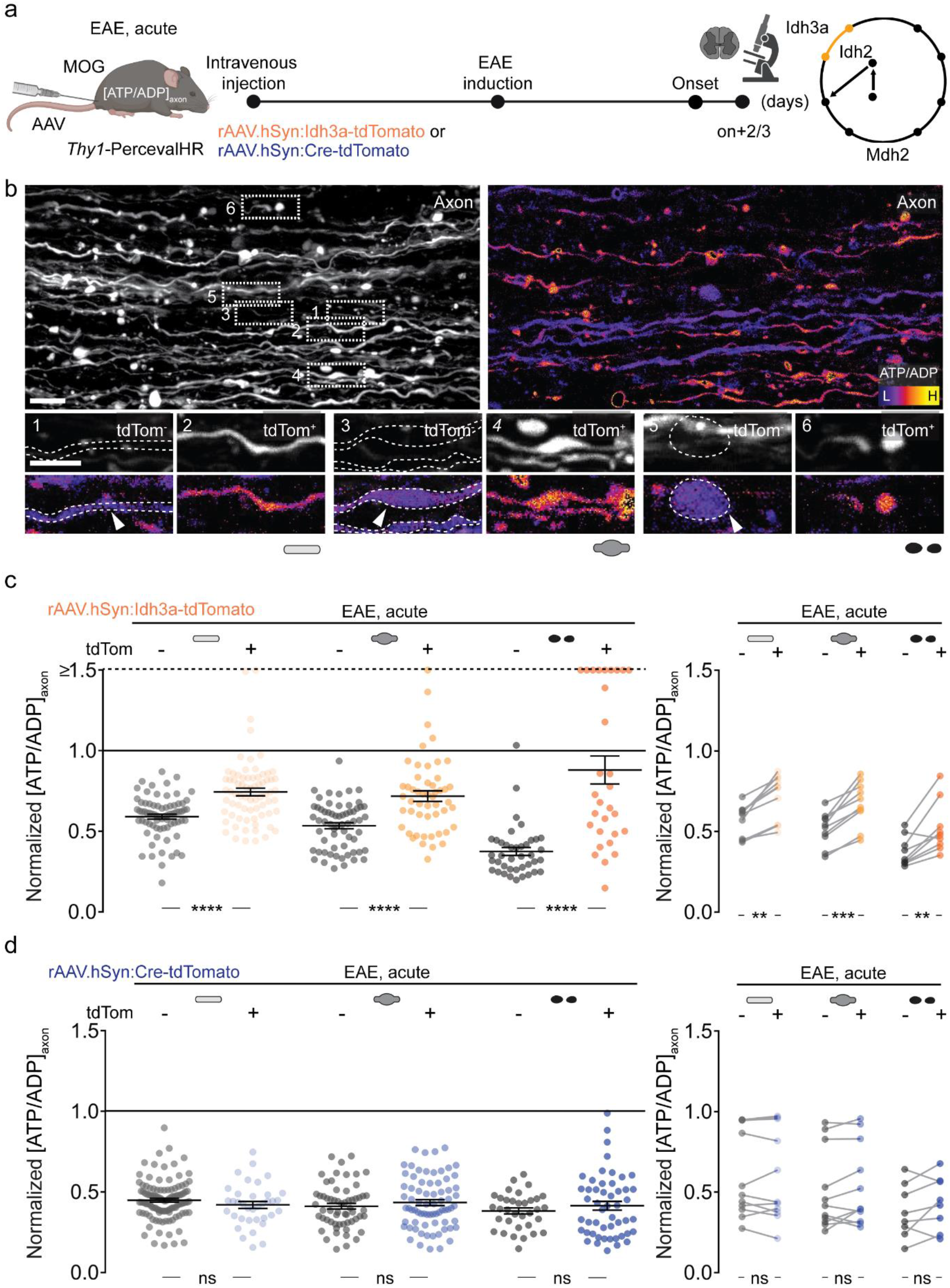
Idh3 overexpression ameliorates axonal ATP deficits in EAE lesions. **(a)** Experimental design for [ATP/ADP]_axon_ in acute EAE in *Thy1*-PercevalHR mice that virally overexpressed Idh3a or a control protein (Cre recombinase) together with tdTomato in a subset of axons. (**b**) Maximum intensity projections of *in vivo* multi-photon image stacks of spinal cord axons in Idh3a-overexpressing *Thy1*-PercevalHR mice. **Left**: Grayscale LUT of tdTomato. **Right**: Ratiometric [ATP/ADP]_axon_ LUT (λ_ex_ ratio 950 nm/840 nm). Details below show image pairs of tdTomato-negative (tdTom^-^, left) and -positive (tdTom^+^, right) normal-appearing, swollen, and fragmented axons (tdTomato-negative with dashed outlines) in acute EAE. (**c**) Comparison of [ATP/ADP]_axon_ in tdTomato-positive and -negative axons (plotted as λ_ex_ ratio 950 nm/840 nm, normalized to control axon mean indicated as the black line; values above 1.5 are lined up on the “≥1.5” dashed line). **Left**: [ATP/ADP]_axon_ of single tdTomato-negative (gray) and -positive (orange) axons in Idh3a-overexpressing EAE mice. **Right**: Lesion-specific paired analysis of mean [ATP/ADP]_axon_ in tdTomato-negative (gray) and -positive (orange) axon populations of the three morphological stages. (**d**) Same analysis as c, but in a mouse cohort overexpressing a control protein (Cre recombinase). Mean ± s.e.m. Comparison of n > 300 axons in > 9 lesions from 4 mice in c; n > 300 axons in > 8 lesions from 3 mice in d using a two-tailed, unpaired Student’s t-test (c, d, left graphs) and a paired t-test (c, d, right graphs). Scale bar: 25 μm in b. **, p < 0.01; ***, p < 0.005; ****, p < 0.001.

Overall, these results suggest that a combinatorial targeting of the TCA cycle, or rather a metabolic approach to boosting TCA cycle function, will be needed to more optimally remedy axonal bioenergetics in neuroinflammation – and indeed, when we explored the effects of Idh3a overexpression on axonal morphologies in tdTomato positive vs. negative axons, we could not detect robust differences in axonal swelling or fragmentation at 3 weeks after disease onset (**Extended Data Figure 6d**).

## DISCUSSION

That axons in neuroinflammatory lesions exist in a state of ‘energy failure’ linked to mitochondrial damage has long been inferred based on (1) abnormalities in mitochondrial density, dynamics and morphology; (2) mitochondrial dysfunction as revealed by in situ assays of respiratory complex function or mitochondrial polarization; and (3) the accumulation of mitochondrial DNA mutations in inflamed CNS tissue^1, 2, 18^. Furthermore, there is molecular evidence that axons in MS lesions might exist in a hypoxic state, which would further restrict mitochondrial respiration^37^. These data have directed most attention to the ETC, which covers most of the uniquely high energy demands of neurons and determines the mitochondrial potential and other aspects of respiration that can be assayed in situ, e.g., using enzymatic assays. Thus, the focus on the ETC is both well justified by its importance, but also biased by experimental accessibility. At the same time, the ETC is not necessarily a good therapeutic target, given its complex molecular make-up at the protein, as well as at the genome level, the extremely low turnover of some of its components and its Janus-faced nature as the major source not only of ATP, but also of oxygen radicals. Finally, the ETC needs to be fed with a steady and sufficient flow of redox equivalents and other substrates, most of which derive from the TCA cycle. Hence the TCA cycle – as the ‘upstream’ bioenergetic hub of metabolism – also deserves attention in the context of the presumed ‘energy failure’ of axons in neuroinflammation^38^. Especially so, as the TCA cycle is increasingly emerging as a metabolic hub relevant to neurodegeneration^25^ and a ‘druggable’ target, for example in cancer^31, 39^.

Against this backdrop, the present study makes the following contributions: (1) We demonstrate directly by *in vivo* biosensor measurements (both using PercevalHR to measure the ATP/ADP ratio, as well as ATeam, which assays ATP levels) that most axons in neuroinflammatory lesion exist in a state of depressed cytoplasmic ATP availability, already during early stages of the axonal degeneration pathway. Moreover, this change precedes overt signs of mitochondrial redox or calcium dyshomeostasis. (2) We show by molecular profiling of neuronal mitochondria not only a disruption of the ETC, but a similarly profound alteration of the ‘up-stream’ TCA cycle, where a number of enzymes, including Idh3 and Mdh2 are depleted in neuronal mitochondria in EAE and MS lesions. (3) Finally, we demonstrate that over-expressing Idh3a (the catalytic subunit of the enzyme) or Mdh2 suffices to increase axonal ATP provision, albeit to a limited extent. This suggests, on the one hand that the axonal energy crisis caused by neuroinflammation is due to a dual deficiency, both of redox substrate provision from the TCA cycle to the ETC, as well of oxidative phosphorylation itself. On the other hand, a full rectification of these neuroenergetic deficits will likely require a manipulation of either a master regulator, or multiple enzyme targets in parallel.

Together, the results we present here expand our understanding of the molecular pathogenesis of immune-mediated mitochondrial damage and propose new avenues for therapeutic intervention. They also raise the question where the dysregulation of the TCA cycle originates. Overall, the pattern of mitochondrial proteome changes does not relate to any simple pattern that we could discern. While the overall mitochondrial mass in neurons cannot be estimated easily by this method, the data suggest that within the mitochondrial proteome of neurons, specific dysregulation happens, as e.g. some biosynthetic pathways (such as mitochondrial translation) are upregulated, while most TCA cycle and ETC components are suppressed. Indeed, the parallel and progressive upregulation of Idh2, while Idh3 is downregulated, suggests a specific metabolic ‘rewiring’ response^40^, rather than just a global loss of mitochondrial biogenesis or increase in mitophagy. Most likely, this response involves biosynthetic, as well as proteostatic mechanisms, as the observed dysregulation of mitochondrial proteins does neither correspond to whether a protein is encoded in the mitochondrial DNA (where mutations are known to accumulate in MS)^2, 8^ vs. in the nucleus, nor with the estimated lifetime^41^. Indeed, the correlation of published neuronal translatome data from a similar model^27^ with our neuronal proteomes provides direct support for an element of ‘anterograde regulation’ from the nucleus, while the long life-times of some down-regulated enzymes (including subunits of Idh3^42^) argue for a parallel degradation, given the swift development of the proteomic phenotype in our acute EAE model (which is present in acute lesions two days after disease onset).

Irrespective of the origin of the reduced ATP availability that our biosensor measurements reveal, the further question arises of what this implies for axonal survival, and hence – given the central role of axon degeneration in MS disease course^43^ – for possible lasting functional consequences. While limited ATP availability is often cited as a sufficient argument to assume axonal demise, and profound and lasting ATP deficiency can clearly cause axon degeneration^44^, our observations rather argue for a chronic ‘dystrophy’ than for an acute bioenergetic collapse of axons in neuroinflammation. Indeed, we find reduced ATP/ADP ratios, ATP levels and TCA cycle enzyme expression in the majority of axons, even those with a normal morphology. From previous work, we know that only a subset of these axons is at immediate risk of degeneration, and some even recover morphological and mitochondrial integrity^6^. Notably, a similarly pervasive pattern of dysfunction has been apparent in our previous studies of axonal transport^17^. As axonal transport is highly ATP dependent, and in converse, mitochondria are a major axonal transport cargo^4^, a vicious cycle could result in the long-term break-down of the ATP supply chain – which might not cause immediate axon degeneration, but rather subacute dysfunction. Furthermore, previous reports of in situ ATP measurements in mouse white matter tracts have shown that neuronal activity puts an immediate strain on energy supply^45^, which could sensitize specific activity-related patterns of degeneration. To explore this notion, but also to better understand the relative importance and metabolic consequences of the two aspects of the axonal energy deficiency emerging from the dual dysregulation of the ETC and the TCA cycle, we also performed metabolic modeling^46^ (see **Methods** for details) based on our analysis of mitochondrial proteomes from control and EAE spinal cords. The modeling predicted a diminished absolute availability of ATP, as well as a reduced ATP/ADP ratio – and also suggested that indeed, with growing ‘metabolic load’ (an increased ATP consumption rate, which in neurons could e.g., relate to increased rates of action potential firing and neurotransmission), these deficits should progress (**Extended Data Figure 8a**). Furthermore, in silico rectification of groups of proteins to their control levels revealed that while ‘normalizing’ the ETC had some effect on ATP levels, the TCA cycle enzymes are predicted to be the more efficacious target point (**Extended Data Figure 8b**).

So how to counterbalance the axonal energy deficiency? Indeed, ‘brain energy rescue’ has been hailed as a possible therapeutic strategy across neurological disorders^47^. The most direct approach would be the provision of energy substrates or key co-factors for the function of bioenergetic enzymes. Indeed, first trials along these lines – e.g. providing biotin – have been conducted in MS, but failed^48^. Notably, biotin is required for the activity of enzymes that feed into the TCA cycle upstream of Idh3 and Mdh2. Our data on TCA cycle disruptions – combined with the previous insights on ETC dysfunction – would argue that simply feeding a substrate into a broken supply chain will indeed not be efficient. Therefore, key bottlenecks in the ATP-providing metabolic pathways need to be unblocked in addition. Our data show, in accordance with our modeling predictions, that while addressing single bottlenecks, such as Idh3 – a key ‘pace-maker’ enzyme of the TCA cycle^29^ – might have some benefits, such a focused intervention would only result in limited changes to the neuronal energy state. While the fact that Idh3 activity can be directly allosterically stimulated by a low ATP/ADP-ratio^49^ and activated by calcium^50^, two changes that are induced in axons in response to an inflammatory challenge, argue that Idh3a overexpression might have a comparably strong unblocking effect on the TCA cycle, our results with Mdh2 overexpression show, that multiple target points exist within this pathway upstream of the TCA cycle. For actual benefits on axonal survival and neurological deficits, we assume that a combinatorial intervention will be needed – in accordance with our observation that even if we extended Idh3 overexpression for several weeks after EAE onset, no clear reduction in axon pathology was apparent. Moreover, we expect the effects of neuroenergetic dystrophy to really only manifest in the slow evolving late states of chronic neuroinflammation that likely drive progression in MS. Unfortunately, animal models of this key phase of MS largely remain elusive. Thus, while the data presented here provide proof of the principle for TCA cycle targeting, they probably do not yet foreshadow a true translational strategy.

Still, the fact that the TCA cycle enzymes are the focus of substantial translational interest^51, 52^, given the key role of some of them (such as Idh1 and Idh2, but also Mdh2) in cancer biology, suggests that a multi-pronged targeting of TCA cycle-related pathways in neuroinflammation might be a viable translational development. Notably, Idh3 appears to be dysregulated in some neoplasms, including glioblastomas^53^. This has resulted in the development of Idh targeting drugs, e.g. for certain leukaemia and brain tumor subtypes^52, 53^. In cancer, the detrimental consequence of Idh mutations appears to be metabolic rerouting that enhances tumor cell proliferation and immune cell dysregulation^34^. Indeed, also in neuroinflammation some aspects of axonal dysfunction resulting from TCA cycle disruptions could be due to metabolite imbalances rather than mere lack of efficient fuelling of the ETC and hence ATP deficiency. In this context, the concomitant upregulation of Idh2 in chronic stages of our model is notable and raises important questions for future exploration, e.g. by metabolomic analysis of cell type-specific mitochondria. Such analysis could be combined with pharmacological intervention, to reroute substrates into the diminished bioenergetic pathways of axons in an inflammatory milieu. For instance, could the available Idh blockers – most of which are deliberately not targeting Idh3 – still be useful in forcing metabolic flow back into the TCA cycle? Similarly, could approaches of increasing mitochondrial mass (such as overexpression of PGC1α, which is beneficial in inflammatory demyelination and regulates Idh3a expression as part of its broad transcriptional effects^11, 52^) be synergistic with targeting the TCA cycle to unblock energy provision in axons and resolve their dystrophy in conditions like MS? The identification of the TCA cycle and some of its key enzymes as a disease contributor and a potential therapy target indeed opens the possibility for such a comprehensive evaluation of strategies to mitigate the neuronal energy crisis in the inflamed CNS.

## ACKNOWLEDGEMENTS

We would like to thank A. Schmalz, J. Schmitt, K. Plesniar, B. Fiedler, Y. Hufnagel and A. Berghofer for excellent technical assistance, D. Matzek, B. Stahr, M. Korica, N. and M. Budak for animal husbandry. We also acknowledge A. Marti Pastor and Y. Hufnagel for help with MitoTag isolations; K. Dyar (Helmholtz Center Munich) for advice on metabolomics, O. Griesbeck (Max Planck Institute of Neurobiology) for constructs, J. Lichtman (Harvard University) for *Thy1*-OFP mice; and C. de la Rosa (BMC/LMU) support for PHP.eB virus establishment.

This project was supported by the Deutsche Forschungsgemeinschaft (DFG) via TRR 274/1 (Projects B03, C02, C05, Z01, Z02 – ID 408885537). Work in M.K.’s laboratory is further financed through grants from the DFG (TRR128, Project B10 and B13), the European Research Council (ERC) under the European Union’s Seventh Framework Program (FP/2007-2013; ERC Grant Agreement n. 310932), the German Federal Ministry of Research and Education (BMBF; Competence Network Multiple Sclerosis) and the “Verein Therapieforschung für MS-Kranke e.V.”. Work in T.M.’s lab is supported by the DFG (CRC870 A11-ID 118803580, Mi 694/8-1, Mi 694/9-1 A03-ID 428663564, FOR Immunostroke) and the ERC under the European Union’s Seventh Framework Program (FP/2007-2013; ERC Grant Agreement n. 616791). T.M. also acknowledges a Pioneer Grant from Doppelganger Biosystem GmbH for metabolic modeling. T.M. and S.L. are members of and supported by the German Center for Neurodegenerative Diseases (DZNE). High resolution microscopy was supported via a DFG instrumentation grant (INST95/1755-1 FUGG, ID 518284373). M.K. and T.M. were further supported by the DFG through a common grant (Ke 774/5-1/Mi 694/7-1) and – together with S.L., F.P. and F.B. –receive support from the Munich Center for Systems Neurology (SyNergy EXC 2145; Project ID 390857198). D.M. is supported by the Swiss National Science Foundation (SNSF; 310030B_201271 & 310030_185321) and the ERC (865026). M.O.B. was recipient of a doctoral fellowship from the Gertrud Reemtsma Foundation and supported by the Emmy Noether Program of the DFG (BR 6153/1-1). G.L. and L.T. were supported by EMBO Fellowships, G.L. further received an SNSF fellowship. Y.T. received support via the TUM Graduate School via the PhD Program ‘Medical Life Sciences and Technology’, I.E. and C.F. via the Munich School of Systemic Neurosciences (GSC 82 – ID 24184143).

## AUTHOR CONTRIBUTIONS

M.K. and T.M. conceived and designed the experiments. Y.T. performed *in vivo* imaging and disease modeling experiments, with contributions from G.L. for mitochondrial calcium and redox imaging. C.F., G.L, D.T., Y.T., R.N., and M.B. established and characterized mouse lines for *in vivo* imaging, L.T., C.F., and Y.T. established, supported and performed the MitoTag analysis, for which S.M. and S.L. did the proteomic analysis. D.E. did the bioinformatics analysis with input from S.M. and S.L. M.Kr., I.W. and D. Me. did histological analysis of MS, while Y.T., I.E. and S.G. did mouse histology and S.M.-L. and D. Ma. contributed the COX IV assay in EAE. A.K. and Y.T. designed and cloned AAV vectors; A.A and F.B. supported to generate the PHP.eB viruses. N.M. and F.P. supported bioenergics analyses. Y.T. coordinated and contributed data analysis across all experimental approaches and designed the final data representation with input from M.K. and T.M. Y.T., M.K. and T.M. wrote the paper with input from all authors.

## ONLINE METHODS

### Animals

All experiments were performed on either postnatal day 3 pups or adult mice according to the protocols on a C57BL/6 (strain designation C57BL/6J, Jackson Laboratories) background at age from 2 to 6 months. All animals were bred and housed under standard conditions with a 12-hours light/12-hours dark cycle. Food and water for mice were provided ad libitum. Female and male were equally allocated into control and experimental groups if not explicitly mentioned otherwise. The experiments were not powered for independent analysis of male and female mice, but separate analysis of female and male mice did not reveal any major sex-specific effects in our analyses of ATP/ADP levels. All experimental procedures were conducted in accordance with regulations of the relevant animal welfare acts and protocols approved by the responsible regulatory office.

To measure neuronal mitochondrial redox changes in spinal axons, we used the *Thy1*-mitoGrx-roGFP mouse line, which selectively expresses the redox sensor Grx-roGFP2 in neuronal mitochondria^19^. To visualize axonal morphology, these mice were crossed to *Thy1*-OFP mice^20^, in which neurons are labelled by cytoplasmic expression of an orange fluorescent protein (OFP). To investigate the proteomic profile of neuronal mitochondria in the context of neuroinflammation, we used the *Gt(ROSA)26Sor* knock-in mouse line MitoTag generated by recombinase-medicated cassette (*loxP*-flanked STOP) exchange into the Rosa26 locus that allows a Cre-dependent expression of GFP targeted to the outer mitochondrial membrane. The MitoTag mouse line is available from The Jackson Laboratory as JAX#032675 (Rosa26-CAG-LSL-GFP-OMM)^23^. *Thy1*-mitoRFP mice, in which tagRFP is localized to the matrix of neuronal mitochondria^19^ were used to ascertain the localization of immunofluorescence signals to neuronal mitochondria.

### Generation of reporter mouse lines

To measure the neuronal ATP-to-ADP ratio, *Thy1*-PercevalHR mouse lines were generated using a blunt-end cloning strategy with Perceval GW1-HR plasmid (Addgene, #49082) purchased from Addgene^13^. In brief, the GW1-Perceval-HR plasmid was digested with Xba1 and EcoRI enzymes, blunted, and inserted into the *Thy1*-vector cut beforehand with XhoI, blunted, and dephosphorylated. After ligation and electroporation, minipreps of ampicillin-selected E. coli cultures were performed, followed by verifying the correct plasmid orientation and correct enzyme digestions. The construct was transfected in human embryonic kidney 293T (HEK) cells, and the cytoplasmic expression was confirmed by confocal microscopy. Maxipreps of the adequate cloning candidates were carried out using Qiagen kits. The Thy1-Perceval-HR plasmid was further linearized using PvuI and EcoRI restrictions enzymes.

To measure the calcium level in neuronal mitochondria, *Thy1*-mitoTwitch2b mouse lines were generated. The Twitch2b-pcDNA3 plasmid^22^ was kindly provided by O. Griesbeck (Max Planck Institute of Neurobiology, Martinsried, Germany). The Twitch2b sensor was extracted using NotI restriction enzyme and inserted into the pCMV:myc-mito plasmid (Addgene, #71542) cut beforehand with NotI restriction enzyme and dephosphorylated. After ligation and electroporation, minipreps of ampicillin-selected E. coli cultures were performed, followed by verifying the correct plasmid orientation and correct enzyme digestions. The construct was transfected in HEK cells and verified by live imaging. After retesting the construct and its functional responsiveness by chemical application in HEK cell culture, maxipreps of the adequate cloning candidates were carried out using Qiagen kits. The pCMV-myc-mito-Twitch2b sequence was further extracted using PmI and XbaI, blunted and inserted into the *Thy1*-vector cut beforehand with XhoI, blunted and dephosphorylated. The *Thy1*-mitoTwitch2b plasmid was linearized using ZraI and AflIII restrictions enzymes. Sequencing of *Thy1*-mitoTwitch2b and *Thy1*-PercevalHR constructs were performed by Eurofins. The sample purity of the linearized DNA was determined using the absorbance ratio at 260 nm vs 280 nm. The concentration of the linearized DNA used for pronuclear injections was 45.2 ng/µl in 60 µl for Thy1-Perceval-HR (with a ratio A260/280 at 1.83) and 36.5 ng/µl in 120 µl (with a ratio A260/280 at 1.83) for *Thy1*-mitoTwitch2b. The generation of the *Thy1*-PercevalHR and *Thy1*-mito-Twitch2b was conducted at the Transgenic Core Facility of the Max Planck Institute for Molecular Cell Biology and Genetics in Dresden by standard pronuclear injections into pseudo-pregnant host mice. For analysis of mitochondrial calcium level in spinal axons *Thy1*-mitoTwitch2b mice were crossed to *Thy1*-OFP mice.

### AAV vector generation and virus production

The AAV.hSyn:Cre plasmid, which encodes for Cre recombinase expression specifically in neuronal populations under the control of the human Synapsin promoter, was generated by replacing the DIO-GFP sequence (Addgene, #50457) to create the AAV.hSyn backbone plasmid. In brief, the AAV.hSyn:DIO-GFP plasmid was initially digested with SalI/XhoI, followed by incubating with BglII to remove DIO.GFP and ligated with GFP.ires.Cre, which was excised from AAV.CMV:GFP.ires.Cre with BglII using Quick Ligase according to the manufacturer’s instruction. Further from AAV.hSyn:GFP.ires.Cre, GFP.ires was excised with BmgBI, and the backbone was ligated to acquire AAV.hSyn:Cre plasmid.

The AAV.hSyn:Idh3a.P2A.tdTomato plasmid, which encodes for Idh3a.P2A.tdTomato expression under the control of the human Synapsin promotor was generated as follows. In brief, the AAV.hSyn:DIO.FGF.P2A.EGFP was initially digested with SalI/HindIII to remove DIO.FGF.P2A.EGFP, ligated with amplified Idh3a fragment (Sino biological, MG53630-UT; forward primer: gtaccggatcctctagaggccgccaccaagc and reverse primer: gcttccgtctaagtctttgactctacgacagatttcttctg) and amplified P2A.tdTomato sequence (forward primer: aaagacttagacggaagcggagccac and reverse primer: tccagaggttgattatcgatacgttacttatacagctcatcca) by using Gibson assembly master mix (New England BioLabs, E2611S). The AAV.hSyn:mMDH2.P2A.tdTomato plasmid, which encodes for Mdh2.P2A.tdTomato expression under the control of the human Synapsin promotor was generated as follows. In brief, the AAV.hSyn:DIO.FGF.P2A.EGFP was initially digested with SalI/HindIII to remove DIO.FGF.P2A.EGFP, ligated with amplified Mdh2 fragment (Sino biological, MG51925-UT; forward primer: gtaccggatcctctagaggccgccaccaagc and reverse primer: ccgcttcccttcatgttcttgacaaagtcctcgcct) and amplified P2A.tdTomato sequence (CAG.Cre.P2A.tdTomato; forward primer: catgaagggaagcggagccactaacttc and reverse primer tccagaggttgattatcgatacgttacttatacagctcatcca), by using Gibson assembly master mix.

The AAV.hSyn:ATeam plasmid, which encodes for ATeam, a FRET-based ATP indicator, under the control of the human Synapsin promoter was generated as follows. In brief, AAV.hSyn:DIO.FGF.P2A.EGFP was initially digested with SalI/HindIII to remove DIO.FGF.P2A.EGFP, ligated with amplified ATeam fragment (Addgene, #51958; forward primer: tccagaggttgattatcgataatagggccctctagatgcatgctc and reverse primer: gtaccggatcctctagaggagacccaagcttggtaccg) by using Gibson assembly master mix.

The AAV.hSyn:SypHer3s plasmid, which encodes for SypHer3s, a ratiometric pH probe, under the control of the human Synapsin promoter was generated as follows. In brief, AAV.hSyn:DIO.FGF.P2A.EGFP was initially digested with SalI/HindIII to remove DIO.FGF.P2A.EGFP, ligated with amplified SypHer3s fragment (Addgene, #108118; forward primer: gtaccggatcctctagaggccgccaccatgtccggacc and reverse primer: tccagaggttgattatcgataccgtcgactgcagaattctca) by using Gibson assembly master mix.

The AAV.hSyn:Cre.P2A.tdTomato plasmid used to express Cre and red fluorescent protein dTomato specifically in neurons was purchased from Addgene (Addgene, #107738). The construct with the correctly oriented insert was introduced to Stellar Competent cells (Clontech, 636763), and the plasmid purification was performed with a Qiagen Plasmid Maxi Kit according to the manufacturer’s protocol.

AAV vector packaging was performed using HEK 293T (ATCC, crl-3216) and produced as described before68. In short, HEK 293T cells were transfected with pAD-helper, AAV-capsid 9 (Addgene, #112865) or PHP.eB^36^ (Addgene, #103005), and the AAV-construct (molar ratio 1:1:1) using a RPMI: PEI incubation protocol. AAV vectors were harvested from the supernatant with polyethylene glycol (PEG, Sigma-Aldrich, no. 25322-68-3) solution and the cell pellets. Freeze-thaw cycles were performed to lyse the cells, and the residual DNA from the packaging was further degraded with benzonase (Sigma-Aldrich, E1014). Following the purification procedure using iodixanol gradient ultracentrifugation, the virus was concentrated by subsequent centrifugation and incubation with formulation buffer (Pluronic-F68 0.001% in saline PBS). The product was then further collected (with a genomic titer of ∼10^12^ to 10^14^) and stored in small aliquots at -80°C.

### Neonatal intraventricular injection of rAAV

For the proteomic profiling of mitochondria in neuroinflammation, AAV.hSyn:Cre was injected into neonatal *MitoTag* pups (genotype: WT/MitoTag) according to a previously published protocol^54^. In short, P3 pups were anaesthetized with isoflurane (Abbott) and injected with 3 μl (∼1 x 10^12^ viral particles per animal) AAV9.hSyn: Cre with 0.05% (wt/vol) trypan blue to the right lateral ventricle at a rate of 30 nl/s using a fine glass pipette (Drummond; 3.5”, #3-000-203-G/X) attached to a nanoliter injector (World Precision Instruments; Micro4 MicroSyringe Pump Controller connected with Nanoliter 2000) held by the stereotaxic manipulator. All surgeries were conducted under ultrasound guidance (VisualSonics; Vevo1100 Imaging System, with a Microscan MS550D transducer).

### Intravenous injection of rAAV

To achieve overexpression of Idh3a or Cre in spinal axons, AAV.PHP.eB.hSyn: Idh3a.P2A.tdTomato, AAV.PHP.eB.hSyn:Mdh2.P2A.tdTomato, AAV.PHP.eB.hSyn:ATeam, AAV.PHP.eB.hSyn:Cre.P2A.tdTomato or AAV.PHP.eB.hSyn:SypHer3s was intravenously injected 14 days prior to the EAE induction. In brief, the animal was immobilized, and the tail was locally warmed to dilate the vessels facilitating the injection. The mice were injected with the virus (∼1 x 10^11^ viral particles per animal) at a final volume of 120 µl, followed by disinfection of the operation area with 80% ethanol.

### Induction of experimental autoimmune encephalitis (EAE)

Mice were immunized subcutaneously with 250 µl of an emulsion containing 200 μg of purified recombinant MOG (N1-125) and complete Freund’s adjuvant (Sigma-Aldrich, F5506) supplemented with 10 mg/ml Mycobacterium tuberculosis (H37RA, BD Difco, 231191). The mice received intraperitoneal (i.p.) injections with 200 ng pertussis toxin at day 0 and 2. After immunization, mice were weighed daily, and neurological deficits were evaluated according to the following EAE score: 0, no clinical signs; 0.5, partial tail weakness; 1, tail paralysis; 1.5, gait instability or impaired righting ability; 2, hind limb paresis; 2.5, hind limb paresis with dragging of one foot; 3, total hind limb paralysis; 3.5, hind limb paralysis and forelimb paresis; 4, hind limb and forelimb paralysis; 5, death. The onset of disease was defined by the first day of neurological symptoms; the acute stage as 2-3 days following clinical onset and the chronic stage as 21/22 days following the onset of EAE in all chronic experiments.

### Spinal cord laminectomy

Mice were imaged at the acute stage of EAE, two days after onset of disease (only animals with an EAE score ≥ 2.5 were included) and at onset plus 22 days for the chronic stage. The mice were anesthetized by intraperitoneal injection of medetomidine (0.5 mg/kg), midazolam (5.0 mg/kg), and fentanyl (0.05 mg/kg) and tracheotomized and intubated to minimize breathing if needed. The dorsal spinal cord was surgically exposed as previously described^6^. In short, the skin was disinfected and incised along the spinal column. The lumbar-dorsal spinal cord was surgically exposed at the position of vertebrates L3 and L4 by performing a laminectomy, and the surgical field was constantly superfused with pre-warmed artificial cerebrospinal fluid (aCSF, 148.2 NaCl, 3.0 KCl, 1.4 CaCl2, 0.8 MgCl2, 0.8 Na2HPO4 and 0.2 NaH2PO4 in mM) to stay moisturized. The vertebral column was then position-fixed on a spinal clamping device (Narishige STS-a), allowing controlled movement in x, y, z directions during the imaging session. A 3.5% agarose well was built up around the spinal opening and filled with aCSF. A standardized imaging protocol was used, with at least 3 images were acquired in volume stacks penetrating up to 100 µm into the tissue from the spinal cord surface (meningeal level). Acute EAE was studied in animals that showed clinical score ≥ 2.5 with confluent lesions present and image regions were defined based on the accumulation of infiltrated immune cells. During the imaging sessions, mice were kept under constant anaesthesia, and their breathing and reflexes were controlled every 30 min. Animals showing signs of traumatic damage after laminectomy were excluded from the analysis.

### *In vivo* multiphoton and confocal imaging

*Imaging of Thy1-PercevalHR mice:* to detect the axonal ATP/ADP ratio, the genetically encoded ATP/ADP indicator PercevalHR was imaged using a set of wavelengths of 950 nm and 840 nm to excite the cpmVenus fluorescent protein sequentially. The signals were collected simultaneously in cyan and yellow channels using emission barrier filter pairs (bandwidth of 455–490 nm and 526–557 nm) on the Olympus MPE-resonant scanner system. Iodoacetic acid (IAA), 10 mM, or Carbonyl cyanide m-chlorophenyl hydrazone (CCCP), 100 μM, at final concentration were added to the imaging solution to obtain the baseline signal of the PercevalHR sensor. For imaging of the effects of Idh3a-tdTomato and Mdh2-tdTomato overexpression on ATP/ADP ratio, we used the same setting as described above. In addition, tdTomato signals were recorded using a wavelength of 1040 and collected in the red channel with a barrier filter pair (bandwidth 655-725nm).

*Imaging of AAV.PHPeB-hSyn:ATeam injected mice:* to detect the axonal ATP level, the genetically encoded ATP sensor, Ateam, was imaged using wavelength of 840 nm to excite mseCFP and cp173-mVenus fluorescent proteins. The emission signals were collected simultaneously in cyan and yellow channels using emission barrier filter pairs (bandwidth of 455–490 nm and 526–557 nm) on the Olympus MPE-resonant scanner system.

*Imaging of Thy1-mitoTwitch2b × Thy1-OFP mice*: to measure mitochondrial calcium levels in axons, the genetically encoded calcium indicator Twitch-2b was excited using a wavelength of 840 nm to excite mCerulean3 and cpVenusCD fluorescent proteins simultaneously^22^. The signals were collected in cyan and yellow channels. To reveal the axonal morphology, the orange fluorescent signal was excited with a wavelength of 750 nm. The customized emission barrier filter pairs with bandwidths of 457-487 nm, 500-540 nm (GaAsP detectors), and 560-600 nm (RXD2 detector) were used on the Olympus MPE-FV1200 system. Images (12 bit) were acquired with a 25x/1.05 dipping cone water-immersion objective, a pixel size of 0.26 mm pixel^-1^ or smaller, a dwell time of 2.0 µs pixel^-1,^ and a laser power of 30 mW measured in the back focal plane. Volume stacks penetrating ∼50 µm into the dorsal spinal cord from the surface were acquired with a Z-spacing of 1 µm.

*Imaging of Thy1-mitoGrx-roGFP × Thy1-OFP mice*: to detect the mitochondrial redox state in spinal axons, the genetically encoded redox indicator Grx-roGFP was sequentially excited at 405 and 488 nm wavelengths as previously described^19^. The signals were collected with emission barrier filter pairs (bandwidth 492-592 nm) in separate channels using a 50/50 beam splitter on an Olympus FV1000 confocal system. Smaller stacks were acquired with a Z-spacing of 1 µm at a frame rate of 0.1-0.2 Hz.

### Image processing and analysis

Images were analyzed with the open-source image analysis software ImageJ (Fiji, http://fiji.sc/Fiji) and Photoshop (Adobe). EAE lesions were identified according to the accumulation of infiltrated immune cells. And the morphology of individual axons was traced and assessed through multi-stacks. To determine PercevalHR, ATeam or SypHer3s signals in axons, background intensities (non-axonal areas) of both channels were measured and subtracted for every axon in every experiment to create a background mask, and pixel-by-pixel ratios were calculated from the mean over three regions of the same axon and further normalized to the mean value of the control mice to eliminate the batch differences. For representative images that do not show intensity variations, maximum intensity projections of image stacks were gamma-adjusted to enhance the visibility of intermediate gray values and processed with a “Despeckle” filter to lower the detector noise using Photoshop software (Adobe). To measure the redox state of a single mitochondrion or clusters of mitochondria signals in *Thy1*-mitoGrx-roGFP × *Thy1*-OFP, the mean intensity values were measured in the 405 and 488 nm channels as previously described^19^. The values for the two channels were divided (405/488 nm) to obtain a ratio related to the redox state of the sensor. This ratio was normalized to the mean value of the control mice. Mitochondrial morphology is quantified as the mitochondrial shape factor calculated by dividing the length by the width of a single mitochondrion as previously described^6^. To measure the calcium levels of a single mitochondrion or clusters of mitochondria in *Thy1*-mitoTwitch2b × *Thy1*-OFP, the mean intensity values were measured in the cyan and yellow channels. The FRET signal (YFP channel) was corrected by subtracting the measured crosstalk-fraction of the CFP signal and determined as cFRET. The mitochondrial calcium levels were then expressed as the background-corrected cFRET/CFP ratio normalized to the mean value of healthy mice as previously established for FRET-based calcium sensors^14^.

### Mitochondrial isolation

Mitochondrial samples for mass spectrometry were prepared from adult male mice described in Fecher at al.^23^ In brief, AAV.hSyn:Cre -injected control and EAE Mito*Tag* mice that contain the GFP-labelled mitochondrial outer membrane in neurons were anesthetized with isoflurane and transcardially perfused with 1× PBS/heparin. The lumbar spinal cord was dissected, weighed, and homogenized with a Dounce glass homogenizer using three complete up-and-down cycles with an A-type pestle in isolation buffer (IB) on ice. The sample was then transferred to a cell disruption vessel and processed with nitrogen cavitation at 800 psi and under stirring at 60 rpm for 10 min. After pressure release, a protease inhibitor (cOmplete™, EDTA-free Protease Inhibitor Cocktail, Sigma-Aldrich, 5056489001) was added to the resulting tissue fraction, and subcellular sediments were removed through two times centrifugation at 600g for 10 min. The resulting post-nuclear tissue fraction was filtered through a 30 μm cell strainer. For immunopurification (IP), the post-nuclear tissue fraction was diluted to a maximal concentration of 2 mg tissue/ mL in immunopurification buffer (IPB) and 50μl microbeads coated with mouse IgG_1_ subtype antibody (Miltenyi Biotec, 130-091-125) against GFP was added to the sample and incubated for 1 hr at 4°C on the shaker. Magnetic-activated cell sorting (MACS) was applied to separate the microbead-coated mitochondria. The LS columns (Miltenyi Biotec, 130-042-401) were placed in a magnetic QuadroMACS^TM^ separator (Miltenyi Biotec, 130-090-976) followed by the equilibration step with 3 mL IPB. The samples were applied to the column in repeating 3ml steps and washed three times with IPB. The columns were removed from the magnetic separator, and the microbead-coated mitochondria were gently flushed out with the plunger. Mitochondria were pelleted by centrifugation at 12,000g for 3 min at 4°C and washed twice with IB (without BSA and EDTA), and the pellets were immediately stored at -20°C for further experiments. Subsequently, the protein amount was determined using the BCA assay (Pierce BCA Protein Assay Kit; Thermo Fisher Scientific, 23227) according to the manufacturer’s instructions. BSA was used as the standard, and the sample buffer was used to correct the measurement alterations caused by detergent or BSA.

### Sample preparation for mass spectrometry

Mitochondria were immunocaptured from the spinal cord according to the above-described protocol. Samples were lysed in a modified RIPA buffer (1% Triton X-100, 0.5% sodium deoxycholate, 0.1% SDS, 150 mM NaCl, 5 mM EDTA, 50 mM Tris–HCl, pH 8). Cell debris and undissolved material were removed by centrifugation at 16,000g for 10 min at 4 °C. Protein concentrations were assessed using the BCA assay. A protein amount of 20 µg was further diluted with distilled water at a 1:2 ratio, and 50 mM MgCl_2_ was added to a final concentration of 10 mM, and DNA/RNA was digested using 12.5U Benzonase. Disulfide bonds were reduced by adding dithiothreitol (DTT) to a final concentration of 10 mM and incubation for 30 min at 37°C. Cysteine alkylation was performed by adding iodoacetamide (IAA) to a final concentration of 40 mM and incubation for 30 min at 24 ℃ in the dark. The alkylation step was quenched by adding 4 µL of 200 mM DTT.

Protein digestion was performed using the single-pot, solid phase, sample preparation (SP3) protocol^55^. Briefly, 10 µL of a 4 µg/µL bead slurry of Sera-Mag SpeedBeads A and B (GE Healthcare) were added to 20 µg of alkylated protein lysate. Protein binding to the magnetic beads was achieved by adding acetonitrile (ACN) to a final volume of 70% (v/v) and mixing at 1200 rpm at 24 °C for 30 min in a Thermomixer (Eppendorf). Magnetic beads were retained in a DynaMag-2 magnetic rack (Thermo Fisher Scientific), and the supernatant was discarded. Detergents were removed using four washing steps with 200 µL 80% (v/v) ethanol. Proteins were digested with 0.25 µg LysC (Promega, V1671) at 37°C for 3 hrs followed by a second digestion step with 0.25 µg trypsin (Promega, V5111) for 16 hr at room temperature. Tubes were placed in a magnetic rack, and peptides were transferred to 0.22 µm Costar Spin-X filter tubes (Corning) to remove the remaining magnetic beads. Samples were dried by vacuum centrifugation. Finally, peptides were dissolved in 20 µL 0.1% (v/v) formic acid (FA). Furthermore, the peptide concentration was estimated using the Qubit protein assay (Thermo Fisher, Q33211).

### Liquid chromatography-tandem mass spectrometry data acquisition

Samples were analyzed on an Easy nLC-1200 nano UHPLC (Thermo Fisher Scientific) coupled online via a Nanospray Flex electrospray ion source (Thermo Fisher Scientific) equipped with a column oven (Sonation) to a Q-Exactive HF mass spectrometer (Thermo Fisher Scientific). An amount of 1.3 µg of peptides was separated on self-packed C18 columns (300mm × 75 µm, ReproSilPur 120 C18-AQ, 1.9 µm; Dr. Maisch) using a binary gradient of water (A) and acetonitrile (B) supplemented with 0.1% formic acid (gradient: 0 min., 2.4% B; 2 min., 4.8% B; 92 min., 24% B; 112 min., 35.2% B; 121 min., 60% B). Full mass spectrometry spectra were acquired at a resolution of 120,000 (automatic gain control (AGC) target: 3E+6). The 15 most intense peptide ions were chosen for fragmentation by higher-energy collisional dissociation (resolution: 15,000, isolation width: 1.6 m/z, AGC target: 1E+5, normalized collision energy (NCE): 26%). A dynamic exclusion of 120 s was applied for fragment ion spectra acquisition. For EAE samples, 2 technical replicates were measured per sample.

### Liquid chromatography-tandem mass spectrometry data analysis

The LC-MS/MS raw data was analysed with the software MaxQuant (Version 1.6.3.3 or 1.6.10.43)^56^. The data was searched against a one gene per protein canonical database of Mus musculus downloaded from UniProt (download date: 2019-03-14, 22294 entries). Carbamidomethylation of cysteines was defined as fixed modification, whereas oxidation of methionines and acetylation of protein N-termini were set as variable modifications. Peptide mass recalibration using the first search option with 20 ppm mass tolerance was enabled. For the main search, peptide and peptide fragment mass tolerances were set to 7 and 20 ppm, respectively. Peptide spectrum match and protein false discovery rates (FDR) were set to 1%, and FDRs were controlled using a forward and reverse concatenated database search approach. The option match between runs was enabled with a matching time window of 1.5 s. Protein label-free quantification was performed based on at least two ratio counts of unique peptides per protein. For EAE, LFQ intensities of technical replicates were averaged. If a protein was detected in < 50 % of the samples within one experimental group (EAE or control), all values were imputed with values drawn from a standard distribution with a mean value which was 2 standard deviations below the mean LFQ value of the entire dataset and a 0.3-fold standard deviation of the overall standard deviation of the data set (to control for zero inflation). The remaining missing values (of proteins detected in > 50 % of the samples within one experimental group) were imputed by multiple imputations by chained equations (MICE, R package ‘mice’). Label-free quantification (LFQ) values were log_2_-transformed. For subsequent analysis, LFQ values were treated as interval scale values. To test the similarity of samples within the respective experimental group and to identify outliers, principal component analysis (PCA) was performed. Subsequently, outliers were removed from further analyses. The first two principal components (PC) with their explained variance of each sample were visualized. To test if a protein was differentially expressed between the EAE and control group, we calculated a Student’s t-test and a fold change of the LFQ values between the experimental groups. To control type I error inflation, p values were corrected according to Bonferroni.

MitoCarta 2.0^57^ was used to annotate mitochondrial proteins as well as their sub-mitochondrial location. Gene set enrichment analysis was performed with the Python package nezzworker (https://github.com/engelsdaniel/nezzworker) with REACTOME (version 7.4) as a reference gene set library. Proteins which are members of gene sets that showed the most extreme normalized enrichment scores (NES) were further analyzed. To test if mitochondrial, non-mitochondrial proteins and proteins from different mitochondrial respiratory chain complexes were equally abundant in EAE and control samples, one-way analysis of variance was calculated to test for statistically significant differences between the mean LFQ values. To assess transcriptional changes of motor neurons in EAE, we re-analyzed previously published data from Schattling et al.^27^, using a custom RDre script with the DESeq2 library. To assess the correlation of protein lifetime and the alterations of neuronal mitochondrial proteome, we extracted the protein half-lives dataset from Fornasiero *et al*^41^. All analysis was done in a R or Python environment (https://github.com/engelsdaniel/mitoproteomics).

### Histology and immunofluorescence staining

Immunofluorescence staining on mouse tissues was performed as described in Fecher et al.^23^ unless noted otherwise. Animals were anesthetized with isoflurane and perfused transcardially with 5000 U/mL heparin (Sigma-Aldrich, H3149) in PBS followed by 4% PFA (Morphisto, 11762.01000) in phosphate buffer (PB) or 4% formalin (Sigma-Aldrich, HT501128). Tissues were kept in 4% PFA or 4% formalin overnight at 4 °C and subsequently cut using a vibratome (Leica). The tissue sections were permeabilized with 0.3% Triton-X100 (Sigma-Aldrich, X100) and blocked using 2% fish gelatine (Sigma-Aldrich, G7765), 2% FBS (Thermo Fisher Scientific, 10500064), and 2% BSA (Sigma, A7030) in PBS for 1 hr. Antigen retrieval procedures were applied to unmask antigens and epitopes. For this purpose, sections were treated with either 10mM citrate buffer pH 6.0, 10 mM EDTA buffer pH 8.0, or citrate-EDTA (10 mM citric acid, 2 mM EDTA and 0.05% Tween20) pH 6.2 for 10 min at RT followed by 30 min at 90°C and 30 min at 4°C. The sections were further rinsed with 0.05% Tween20 (Sigma-Aldrich, P9416) before the blocking procedure. Primary antibodies were incubated overnight at 4°C at a dilution of 1:400 (Mdh2, Novus Biologicals, NBP1-32259; Idh3a, Novus Biologicals, NBP1-32396; Idh2, Thermo Scientific, 702713) or 1:1000 (RFP, Novus Biologicals, NBP1-97371 and GFP, Abcam, ab13970). The sections were incubated with secondary antibodies at a dilution of 1:1000 for 2 hrs and mounted using Vectashield mounting medium (Vector Laboratories, H-1000) on the following day.

Heat-induced antigen retrieval was performed on PFA fixed tissue following deparaffination for human tissue sections. After 10% FCS/PBS unspecific binding blockade, sections were incubated with primary antibodies in Dako Diluent (Dako, 52022) overnight at 4°C. After washing with Wash Buffer (Dako, 53006) and autofluorescence removal treatment (Merck, 2160), secondary antibodies and DAPI (Life, D3571) at a dilution of 1:200 were incubated at RT for 1 hr and further mounted with Fluoromount Mounting Medium (Sigma, F4680). The use of human samples followed institutional ethical guidelines and was approved by the ethics committee of the University of Geneva (Switzerland). Written informed consent to use autopsy samples for research purposes was obtained for all samples, with the exceptions of autopsies that were performed more than 20 years ago. In all cases, no samples were used from patients who refused involvement in research projects. Patient information is provided in **Extended Data Table 2**. Data from the human cortex are representative of two experiments performed on 7 different cases.

### Confocal imaging and image processing

Sections stained by immunofluorescence labelling to quantify mitochondrial proteins in neuronal somata, were imaged with an upright Olympus FV1000 confocal system equipped with x10/0.4 air, x20/0.85, and x60/1.42 oil immersion objectives or Leica SP8 equipped with ×20/0.75 HC PL APO CS2 and ×40/1.30 oil immersion HC PL APO CS2 objectives. Images were obtained using standard filter sets and processed with Fiji. For representative figure panels, different channels of image series were pseudo-color-coded in Fiji or Adobe Photoshop; schematics were created using BioRender.com. Contrast and brightness were equally adjusted across the entire image. For the panels displayed in **Figure 4** for intensity comparison, immunofluorescence images of both control and EAE *Thy1*-mitoRFP tissues were acquired with the same settings and were adjusted with the same processing parameters. In panels that do not primarily illustrate quantitative differences, gamma was adjusted nonlinearly to enhance the visibility of low-intensity objects. Figures were assembled in Adobe Illustrator.

Human tissue sections were scanned using a whole slide scanner (Pannoramic P250 II, 3DHistech). Regions of interest were manually selected using SlideViewer software (v2.3, 3DHistech) and exported as individual images for further processing using a custom rule-based script in Definiens Developer XD (v2.7, Definiens AG). In short, specific signals for each marker were detected based on their respective intensity and ratio to tissue background intensity. NF70 (neurofilament) was detected first, and Idh3a detection was restricted to the inside of NF70 positive structures. The total area was quantified for each marker. Additionally, mean intensities were exported for each object individually. For quantification of TCA cycle enzymes in mouse axons, we followed the same approach both for reasons of consistency, but also because mitochondria in axons can be difficult to stain in non-paraffin embedded tissue. For this, slides of *Thy1*-MitoRFP mice stained with NF70 together with the mitochondrial markers Mhd2, Idh2 or Idh3a were imaged on Leica SP8 with ×40/1.30 oil immersion using standard filter sets and processed with Fiji. Notably, RFP fluorescence is abolished in this protocol, so independent masking of mitochondria based on a mitochondrial transgene or staging of FAD were not possible in this data set. Image analysis was further performed in Visiopharm (version 2021.09). In summary, NF70-positive structures were detected and taken as the reference regions. Mitochondrial marker-positive puncta were then detected by a 120% intensity cut-off and a minimum size 4 pixels cut-off and quantified strictly within NF70-positive structures. Total area was quantified for NF-positive structures and marker positive puncta, with the ratio representing “occupancy”. Within the thus defined mitochondrial voxel, the mean intensity value for each individual object was quantified. Data was processed using R (version 4.2.2, R-project.org).

### Sequential cytochrome c oxidase (COX) histochemistry and immunofluorescence histochemistry

To assess COX activity in single axons, COX histochemistry was combined with immunofluorescence histochemistry with a primary antibody against the orange fluorescent protein (OFP) followed by a directly conjugated secondary antibody. This sequential technique was performed in the same tissue section and has already been described and validated in the previous studies^5, 24, 58^. COX media consisted of 100μM cytochrome c, 4mM diaminobenzidine tetrahydrochloride, and 20μg/ml catalase in 0.1M phosphate buffer pH7.0. Cryosections were incubated at 37 °C for 30 min and washed in PBS. The cryosections were processed through the immunofluorescent histochemistry steps as mentioned above. The sequentially stained sections were mounted in glycerol with Hoechst nuclear stain and stored at -20 ⁰C until required for imaging by the Zeiss Imager Z1 Apotome 2 microscope.

To assess COX activity in axons, a mask of the individual axon identified by the fluorescent labelling of OFP was generated in Image J and superimposed onto the brightfield image of COX histochemistry. The total area occupied by COX active elements within a single axon was determined as a percentage of the axonal area. Twenty axons (at least 25μm in length) per region were randomly selected from each animal for quantitation. Assessors were blinded by coding the axons in ascending numerical order.

### Metabolic modeling

Metabolic modeling was performed using the QSM™ data analysis platform provided by Doppelganger Biosystem GmbH, using the mass spectrometry analysis described above for *MitoTag*-derived neuronal mitochondria from acute (2-3 days following clinical onset) EAE vs. controls (n = 6, 5 mice respectively). A detailed description of the used approach can be found in Berndt et al.^46^. In brief, the QSM™ kinetic model includes the major cellular metabolic pathways of mitochondrial energy metabolism and glycolysis, as well as key electrophysiological processes at the inner mitochondrial membrane (membrane transport of various ions, mitochondrial membrane potential, generation and utilization of the proton motive force). First-order differential equations govern the time-dependent variations of model variables (i.e. concentration of metabolites and ions), with time-variations of small ions modelled by kinetic equations of the Goldman-Hodgkin-Katz type. The model uses numerical values for kinetic parameters of the enzymatic rate laws from reported kinetic studies of the isolated enzyme, while maximal enzyme activities (*v_max_* values) are estimated based on functional characteristics and metabolite concentrations of a health neuronal tissue (see Berndt et al.^46^). To establish individual metabolic models, the approach uses the protein intensity profiles of quantitative shotgun proteomics of *MitoTag*-derived neuronal mitochondria. The maximal activities of enzymes and transporters are scaled, exploiting the fact that the maximal activity of an enzyme is proportional to the abundance of the enzyme protein via *v_max_ (EAE) = v_max_(normal)*(E(EAE)/E(mean controls))*. *“v _max_(normal)*” denotes the maximal activity for a given enzyme as used in Berndt et al.^46^, while “*E(EAE)*” denotes the protein abundance measured of enzyme E in a given EAE sample and “*E(mean control)*” stands for the mean protein abundance of an enzyme E averaged across the control sample measurements. Applying the model involves a tailored quality control, which evaluates both the number of proteins of interest found (“QC score”) and the number of metabolic processes associated with the enzymes found (“QSM score”), which indicated good suitability for applying the QSM™ kinetic model (QC >90%; QSM>100% with 75% being the standard cut-off). To evaluate energetic capacities, the model calculates the changes of metabolic state due to increasing the rate of ATP consumption above the resting value, with the ATP consumption rate being modelled by a generic hyperbolic rate law of the form *v_*ATP*_*K_*load*\*_(*ATP/ATP+K_m_*). To model increased “Metabolic load”, the parameter “*K_load_* ” was increased in steps until convergence of the ATP production rate to its maximal value.

### Statistics

Statistical analysis was performed using Microsoft excel software and GraphPad Prism (GraphPad Software, Version 7.0, La Jolla, USA). Sample sizes were chosen according to previous *in vivo* imaging studies of spinal axons^6, 14^. Normality of distribution was assessed with the Shapiro-Wilk test. When normal distribution was confirmed, a Student’s t-test was applied for two groups comparison, and one-way ANOVA followed by Turkey’s post hoc comparison was used for more than two groups. The Kruskal–Wallis test followed by Dunn’s multiple comparisons test or Mann–Whitney U test was used where normal distribution could not be confirmed. Obtained p-values were stated as significance levels in the figure legends (**** P<0.0001, *** P<0.001; **P<0.01; *P<0.05).

## SUPPLEMENTARY INFORMATION

**Extended Data Table 1: Fold changes of all mitochondrial proteins detected by MitoTag proteomics in EAE vs. Control spinal cord.**

Uploaded as separate excel sheet

**Extended Data Table 2:** Information on human tissue samples used in this study.

| Case | Age | Sex | Histological specimen | State |
| --- | --- | --- | --- | --- |
| MS1 | 41 | female | Frontal lobe, subcortical lesion | Chronic active |
| MS3 | 45 | female | Frontal lobe, subcortical lesion | Chronic active |
| MS4 | 51 | female | Right frontal-parietal -lobe | Chronic active |
| MS5 | 55 | male | Left frontal lobe | Chronic active |
| MS6 | 58 | male | Frontal lobe, subcortical lesion | Chronic |
| MS8 | 25 | female | Frontal lobe, subcortical lesion | Acute active |
| MS10 | 77 | female | Not indicated | Acute active |

