## Supplementary information for "Targeting the TCA cycle can ameliorate widespread axonal energy deficiency in neuroinflammatory lesions"

### EXENDED DATA FIGURES

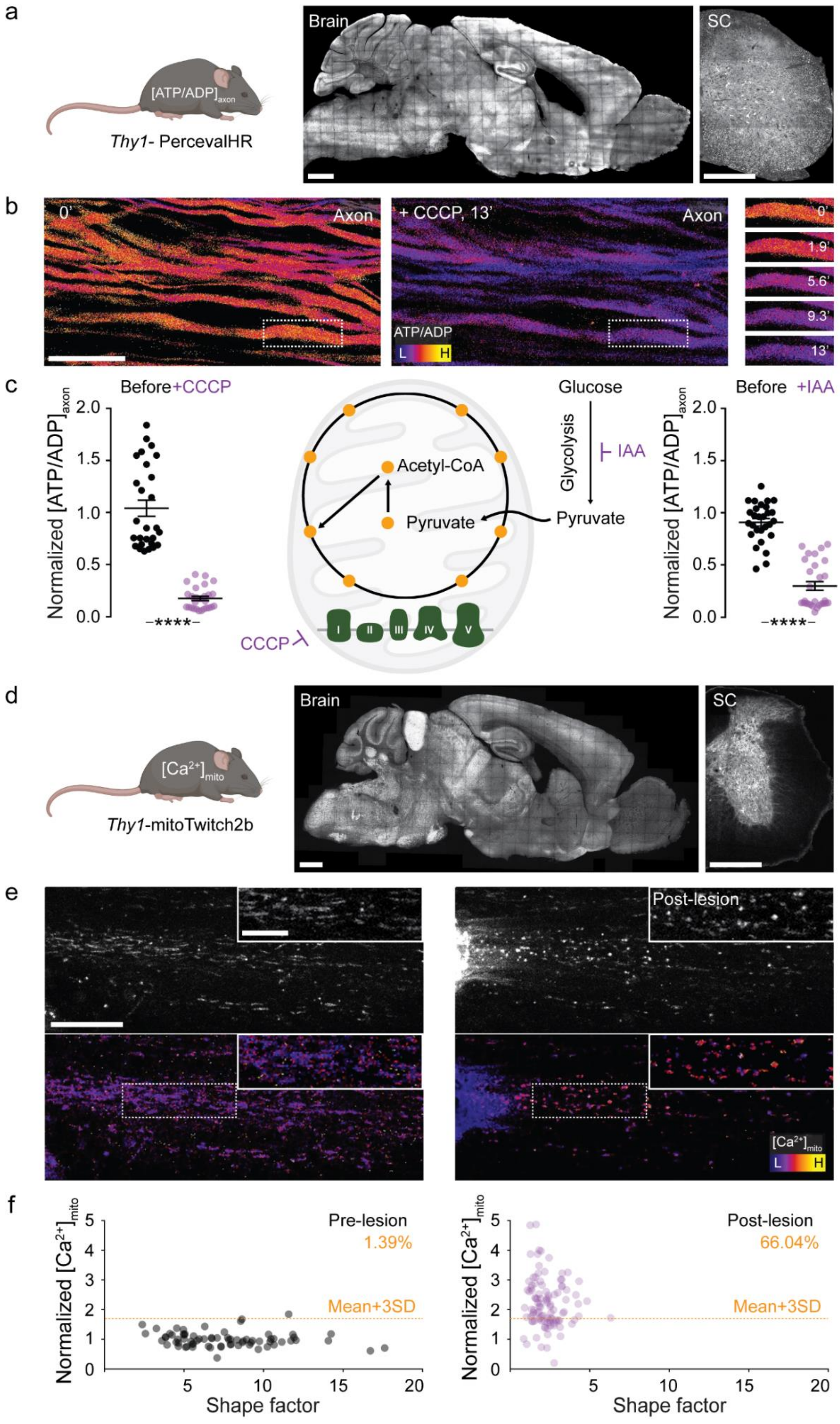


**Tai et al., Extended Data Figure 1**

**Extended Data Figure 1: Transgenic mouse lines to measure neuronal ATP/ADP ratio and mitochondrial calcium.**

(**a**) CNS expression pattern of *Thy1*-PercevalHR mice. Confocal images of sagittal brain (left) and transverse spinal cord (SC) section (right; grayscale LUT; $\lambda_{ex}$ 488 nm). (**b**) Maximum intensity projections of *in vivo* multi-photon image stacks of axons in a *Thy1*-PercevalHR mouse spinal cord before (left) and after (middle) CCCP application. Individual frames in boxed area spanning 0 to 13 minutes after CCCP application (right). Ratiometric $[ATP/ADP]_{axon}$ LUT ($\lambda_{ex}$ ratio 950 nm/840 nm). (**c**) $[ATP/ADP]_{axon}$ of single axons before and after CCCP (100 µM, 15 min; left) and IAA (10 mM, 15 min; right) application, which interfere with oxidative phosphorylation and glycolysis, respectively. Values are normalized to mean of control (mean ± s.e.m.; n > 25 axons, 3 mice/condition compared by Student's t-test). Scale bars: 1000 µm in a, left; 500 µm in a, right; 25 µm in b. ****, $p < 0.001$. (**d**) CNS expression pattern of *Thy1*-mitoTwitch2b mice. Confocal images of sagittal brain (left) and transverse SC section (right; YFP channel using grayscale LUT). (**e**) Maximum intensity projections of *in vivo* multi-photon image stacks of axonal mitochondria in a *Thy1*-mitoTwitch2b mouse spinal cord before (left) and after (right) laser lesion. Top: YFP channel using grayscale LUT; Bottom: Ratiometric $[Ca^{2+}]_{mito}$ LUT (YFP/CFP emission ratio). Insets: Details from respective panels. (**f**) $[Ca^{2+}]_{mito}$ of axonal mitochondria before and after laser lesion (n > 150 mitochondria from 8 axons, 3 mice). Percentages indicate the fraction of axons with $[Ca^{2+}]_{mito}$ > mean + 3SD of values pre-lesion (orange line). Scale bars: 1000 µm in d, left; 500 µm in d, right; 20 µm in e; 10 µm in inset.

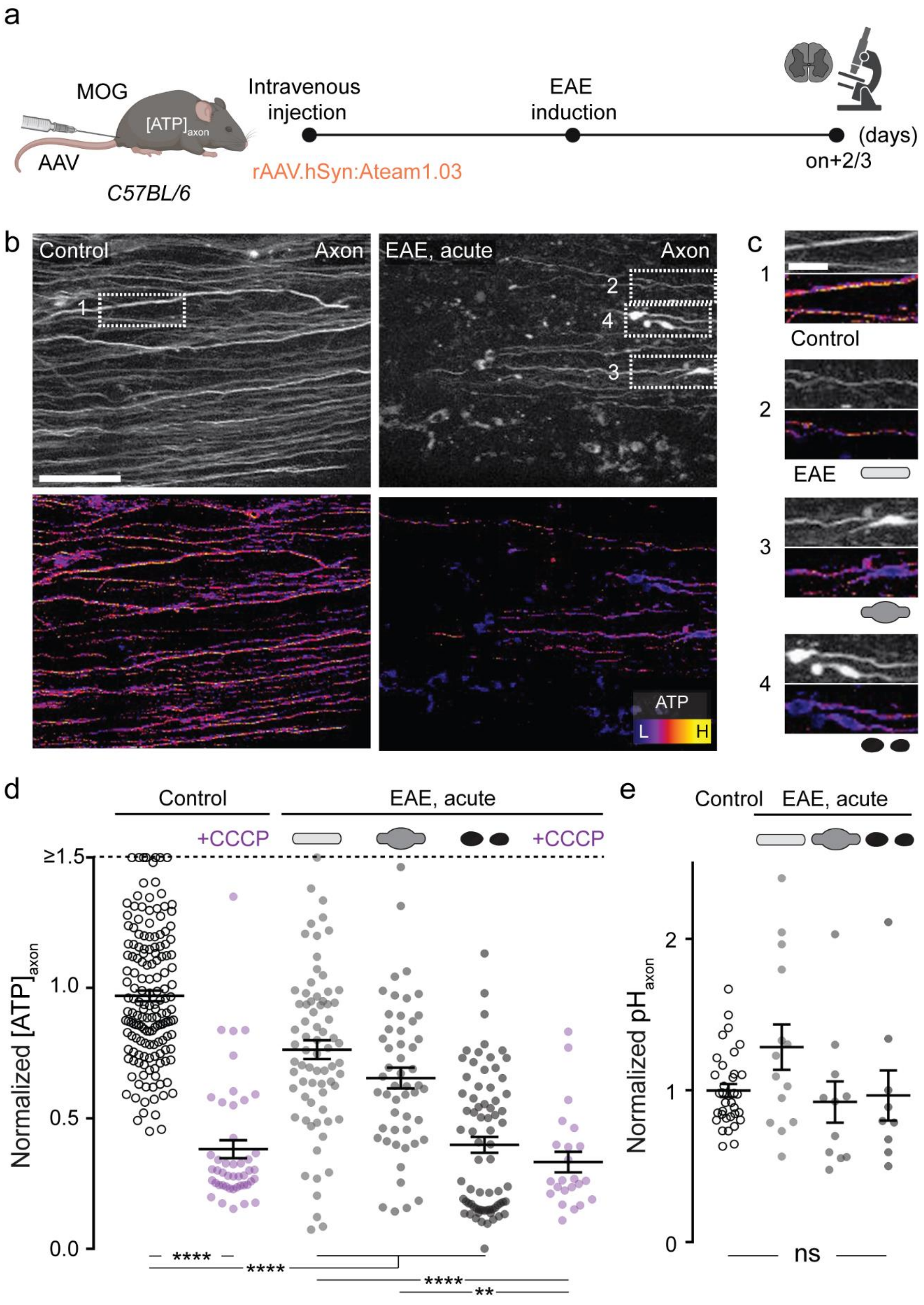


**Tai et al., Extended Data Figure 2**

**Extended Data Figure 2: Early axonal ATP deficits measured using ATeam in EAE.**

(**a**) Experimental design for axonal ATP level measurement in experimental autoimmune encephalomyelitis (EAE). (**b**) Maximum intensity projections of *in vivo* multi-photon image stacks of spinal cord axons of control (left) and acute EAE (right) in AAV.PHPeB.hSyn:ATeam injected C57BL/6 mice. Top: Grayscale look-up table (LUT; $\lambda_{ex}$ 840 nm). Bottom: Ratiometric $[ATP]_{axon}$ LUT (YFP/CFP emission ratio). (**c**) Details from b. Top to bottom: $[ATP]_{axon}$ images of control axon in healthy spinal cord, and normal-appearing, swollen, and fragmented axons in acute EAE. (**d**) $[ATP]_{axon}$ of single axons in healthy and EAE mice normalized to mean of controls (mean ± s.e.m.; $n > 400$ axons, 3 control and 4 EAE compared by Kruskal-Wallis and Dunn's multiple comparison test; values above 1.5 lined up on the "≥1.5" dashed line). (**e**) $[pH]_{axon}$ of single axons measured by using SypHer3s sensor in control and acute EAE mice normalized to mean of controls. (mean ± s.e.m.; $n > 60$ axons, 2 control and 2 EAE mice compared Kruskal-Wallis and Dunn's multiple comparison test). Scale bars: 25 μm in b; 10 μm in c. **, $p < 0.01$; ****, $p < 0.001$.

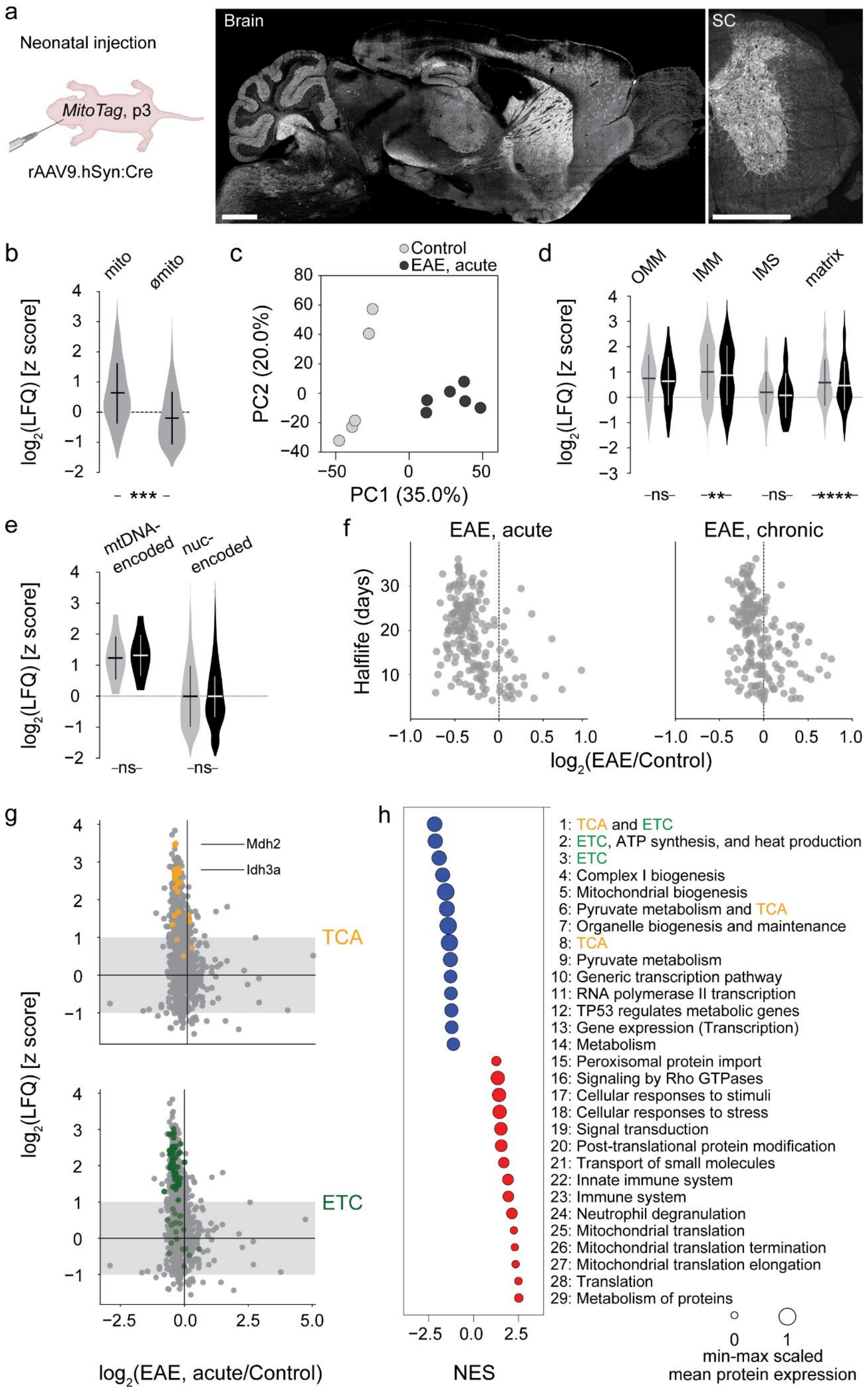


**Tai et al., Extended Data Figure 3**

**Extended Data Figure 3: Proteomics analysis of neuronal mitochondria in EAE.**

(**a**) CNS expression pattern of *MitoTag* mouse injected with rAAV.hSyn:Cre. Confocal images of the sagittal brain (left) and transverse spinal cord (SC) section (right; grayscale LUT; $\lambda_{ex}$ 488 nm). (**b**) Relative expression level (z-score for label-free quantification intensity, LFQ) of proteins quantified by mass spectrometry in *MitoTag* isolations that are annotated as mitochondrial proteins in MitoCarta 2.0[57] (mito) or not (ϕmito). (**c**) Principal component analysis of MitoTag proteomics control and acute EAE samples. (**d**) Expression level ($\log_2$ of LFQ) of proteins annotated in MitoCarta 2.0[57] to reside in different mitochondrial subcompartments in control (gray) and acute EAE (black), including outer membrane (OMM), inner membrane (IMM), intermembrane space (IMS) and matrix relative to all MitoCarta proteins. (**e**) Same as d, but for nuclear- vs. mitochondrial DNA-encoded proteins. (**f**) Fold change ($\log_2$ of LFQ ratio of EAE over Control) versus protein half-life (as measured in Fornasiero *et al*[41]). (**g**) Location of the TCA cycle (top, orange) and electron transport chain (ETC; bottom, green) proteins on the correlation plot of protein abundance (z score of label-free quantification values, LFQ) vs. fold change ($\log_2$ of LFQ ratio acute EAE over control). Gray area, mean ± 1SD. Biological replicates: n≥3. (**h**) Annotations of the most regulated pathways (Reactome[59,60], version 7.4) in *MitoTag* proteomes of neuronal mitochondria in acute EAE. Scale bars: 1000 μm in a, left; 500 μm in a, right.

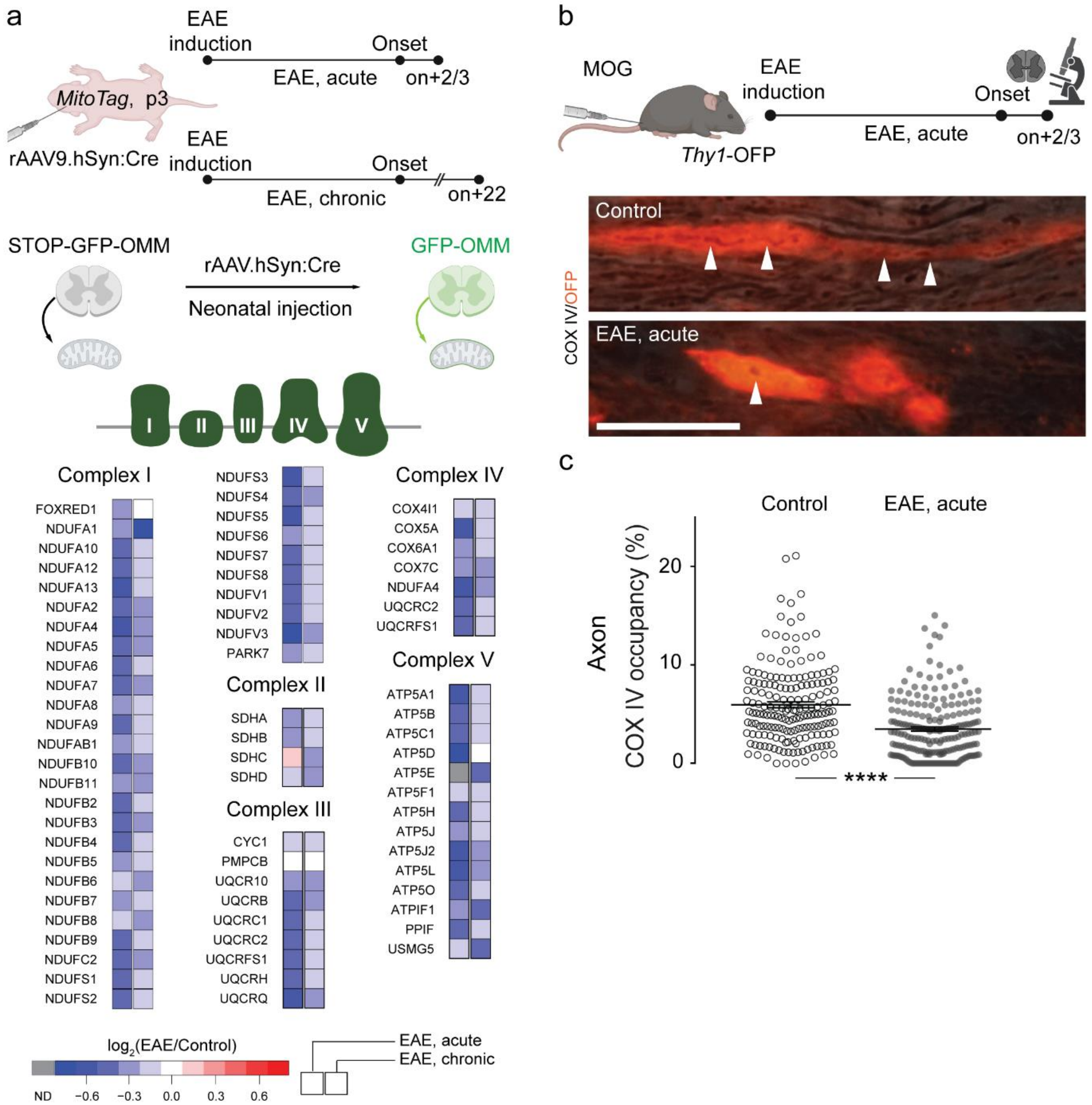


**Tai et al., Extended Data Figure 4**

**Extended Data Figure 4: Expression of ETC components in EAE neurons.**

(**a**) Top: Schematic of the experiment, analysis of same data sets as shown in Fig. 3. Bottom: Relative abundance of individual ETC complex components in neuronal mitochondria. Average shown as color-coded $log_2$(EAE/Control) for acute and chronic EAE compared to respective controls. (**b**) *In situ* analysis of axonal COX IV activity using *in situ* histochemical assay in *Thy1*-OFP EAE spinal cords. Top: Schematic of the experiment. Bottom: Confocal image of control and EAE stage 1 axons, OFP (red) and COX IV (arrow heads indicate mitochondria as dark areas, as fluorescence of OFP is quenched by reaction product of COX IV assay). (**c**) Quantification of COX IV activity signal's occupancy of OFP axon area on axonal level (mean ± s.e.m.; n > 350 axons, 9 mice for control and EAE each, tested with unpaired t test). Scale bar: 25 μm in b. **, $p < 0.01$; ****, $p < 0.001$.

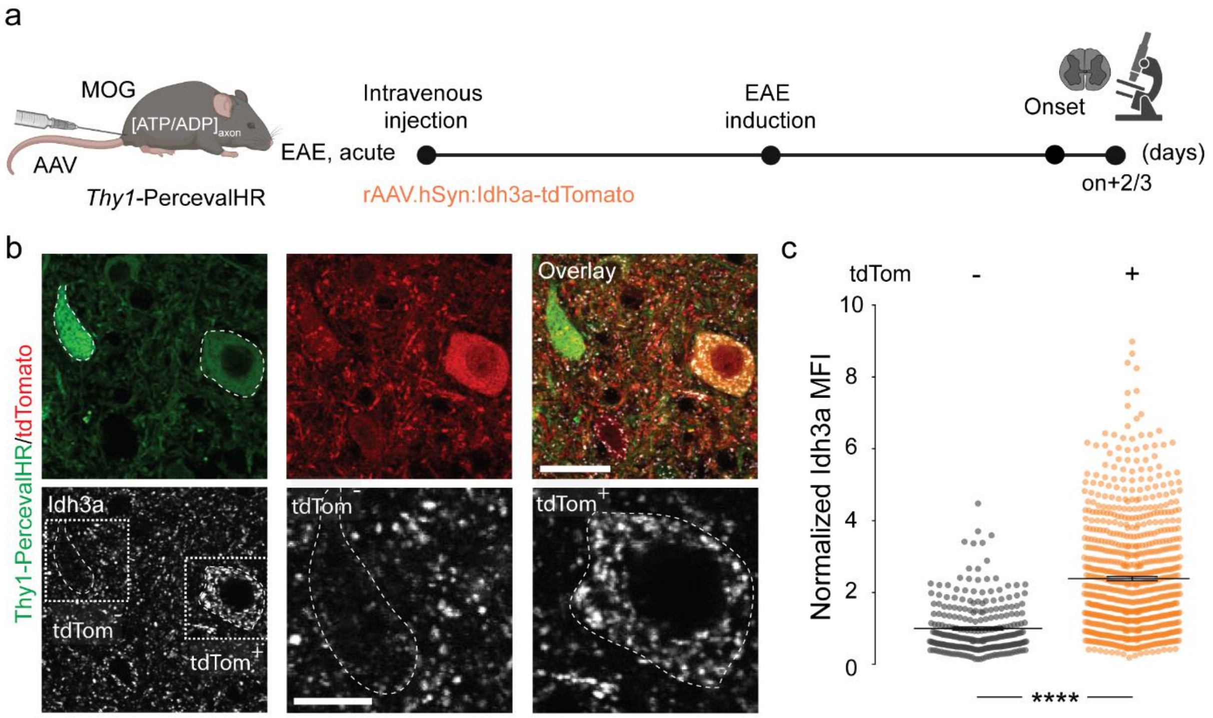


**Tai et al., Extended Data Figure 5**

**Extended Data Figure 5: Viral overexpression of Idh3a in EAE neurons.**

(**a**) Schematic of experiment, analysis of same data sets as shown in Fig. 6. (**b**) Confocal images of spinal cord sections of a *Thy1*-PercevalHR mouse (green) that was injected with rAAV.hSyn:Idh3a-tdTomato (tdTomato, tdTom, red). Bottom row shows immunostainings for Idh3a, with details highlighting, left, a tdTomato-negative ($tdTom^-$) and, right, a tdTomato-positive ($tdTom^+$) neuron with Idh3a overexpression. (**c**) Expression level of Idh3a in tdTomato-negative ($tdTom^-$) and tdTomato-positive ($tdTom^+$) neurons (mean ± s.e.m.; n > 1000 neurons in 20 sections, unpaired t-test). Scale bar: 25 µm (top) and 10 µm (bottom) in b. ****, p < 0.001.

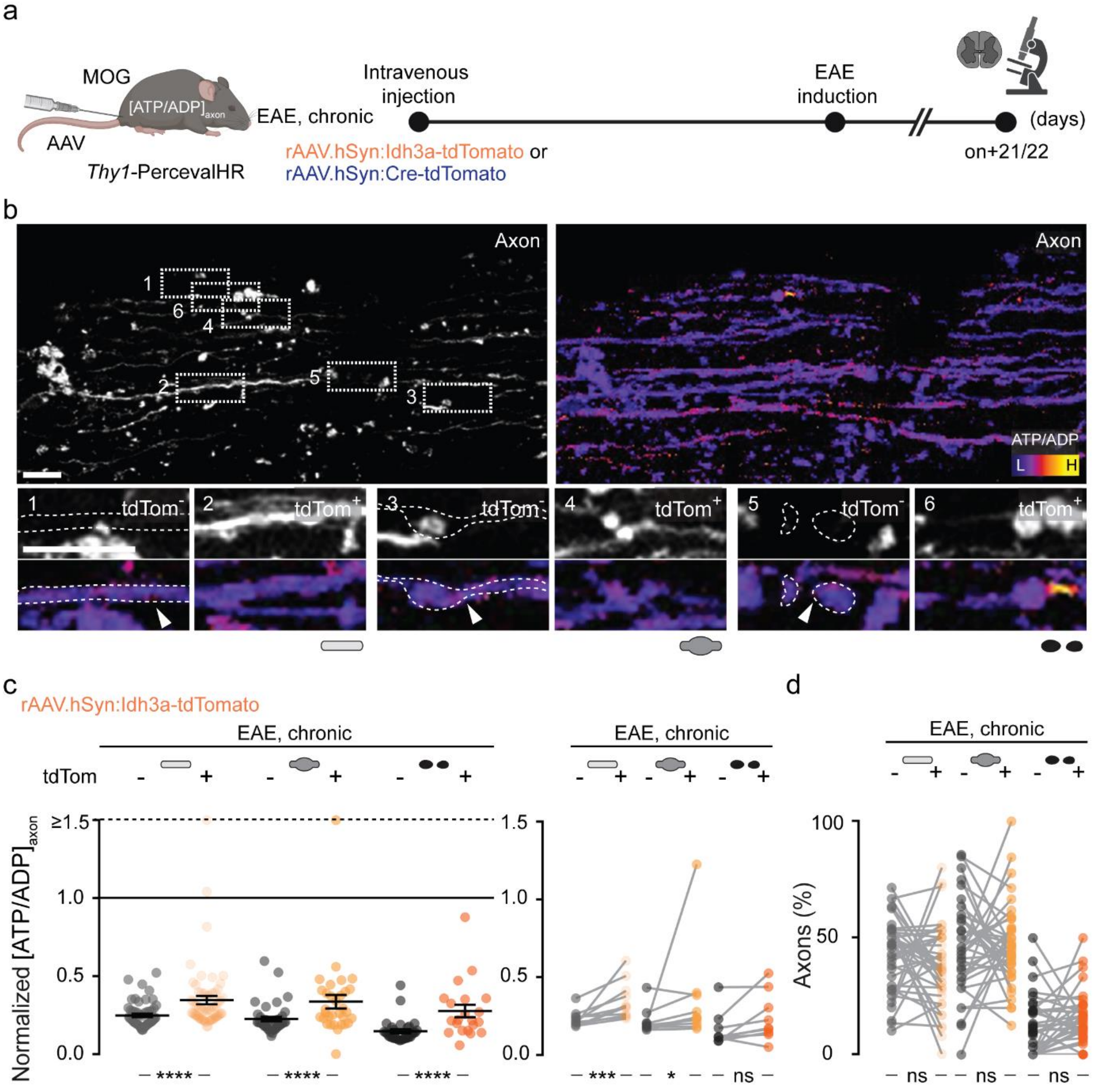


Tai et al., Extended Data Figure 6

**Extended Data Figure 6: Chronic Idh3 overexpression modestly ameliorates axonal ATP deficits in EAE lesions.**

(a) Experimental design for $[ATP/ADP]_{axon}$ measurements in chronic EAE in *Thy1*-PercevalHR mice that virally overexpressed Idh3a or a control protein (Cre recombinase) together with tdTomato in a subset of axons. (b) Maximum intensity projections of *in vivo* multi-photon image stacks of spinal cord axons in chronically Idh3a-overexpressing *Thy1*-PercevalHR mice. Left: Grayscale LUT of tdTomato. Right: Ratiometric $[ATP/ADP]_{axon}$ LUT ($\lambda_{ex}$ ratio 950 nm/840 nm). Details show image pairs of tdTomato-negative (left) and -positive (right) normal-appearing, swollen, and fragmented axons in chronic EAE. (c) Comparison of $[ATP/ADP]_{axon}$ in tdTomato-positive and -negative axons (plotted as $\lambda_{ex}$ ratio 950 nm/840 nm, normalized to control axon mean indicated as the black line; values above 1.5 lined up on the "≥1.5" dashed line). Left: $[ATP/ADP]_{axon}$ of single tdTomato negative (gray) and positive (orange) axons in Idh3a-overexpressing EAE mice. Right: Lesion-specific paired analysis of mean $[ATP/ADP]_{axon}$ in tdTomato-negative (gray) and -positive (orange) axon populations of the three morphological stages. Mean ± s.e.m. Comparison of n > 200 axons in > 9 lesions from 4 mice using a two-tailed, unpaired Student's t-test (left graph) and a paired t-test (right graph). (d) Paired analysis of the frequency of stage 0, 1 and 2 axons in tdTomato-negative (gray) and -positive (orange) axon populations. Mean ± s.e.m. Comparison of n > 35 lesions from 4 mice in d using a paired t test. Scale bars: 25 μm in b. *, $p < 0.05$; ***, $p < 0.005$; ****, $p < 0.001$.

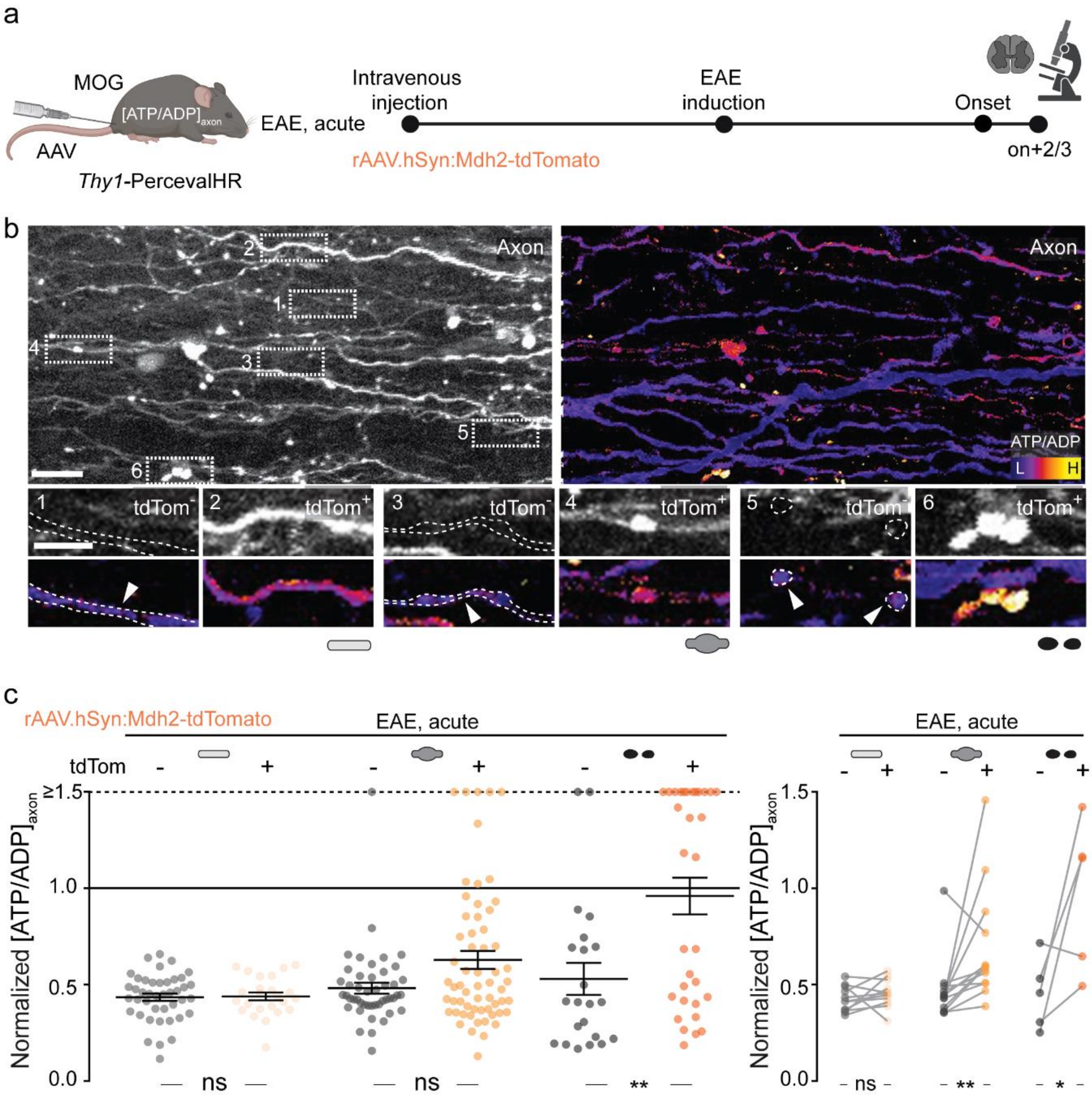


**Tai et al., Extended Data Figure 7**

**Extended Data Figure 7: Mdh2 overexpression ameliorates axonal ATP deficits in EAE lesions.**

(**a**) Experimental design for $[ATP/ADP]_{axon}$ in acute EAE in *Thy1*-PercevalHR mice that virally overexpressed Mdh2 together with tdTomato in a subset of axons. (**b**) Maximum intensity projections of *in vivo* multi-photon image stacks of spinal cord axons in Mdh2-overexpressing *Thy1*-PercevalHR mice. Left: Grayscale LUT of tdTomato. Right: Ratiometric $[ATP/ADP]_{axon}$ LUT ($\lambda_{ex}$ ratio 950 nm/840 nm). Details show image pairs of tdTomato-negative (left) and -positive (right) normal-appearing, swollen, and fragmented axons in acute EAE. (**c**) Comparison of $[ATP/ADP]_{axon}$ in tdTomato-positive and -negative axons (plotted as $\lambda_{ex}$ ratio 950 nm/840 nm, normalized to control axon mean indicated as the black line; values above 1.5 lined up on the "≥1.5" dashed line). Left: $[ATP/ADP]_{axon}$ of single tdTomato negative (gray) and positive (orange) axons in Mdh2-overexpressing EAE mice. Right: Lesion-specific paired analysis of mean $[ATP/ADP]_{axon}$ in tdTomato-negative (gray) and -positive (orange) axon populations of the three morphological stages. Mean ± s.e.m. Comparison of n > 200 axons in > 9 lesions from 4 mice using a two-tailed, unpaired Student's t-test (left graph) and a paired t-test (right graph). Scale bar: 25 µm in b. *, $p < 0.05$; **, $p < 0.01$.

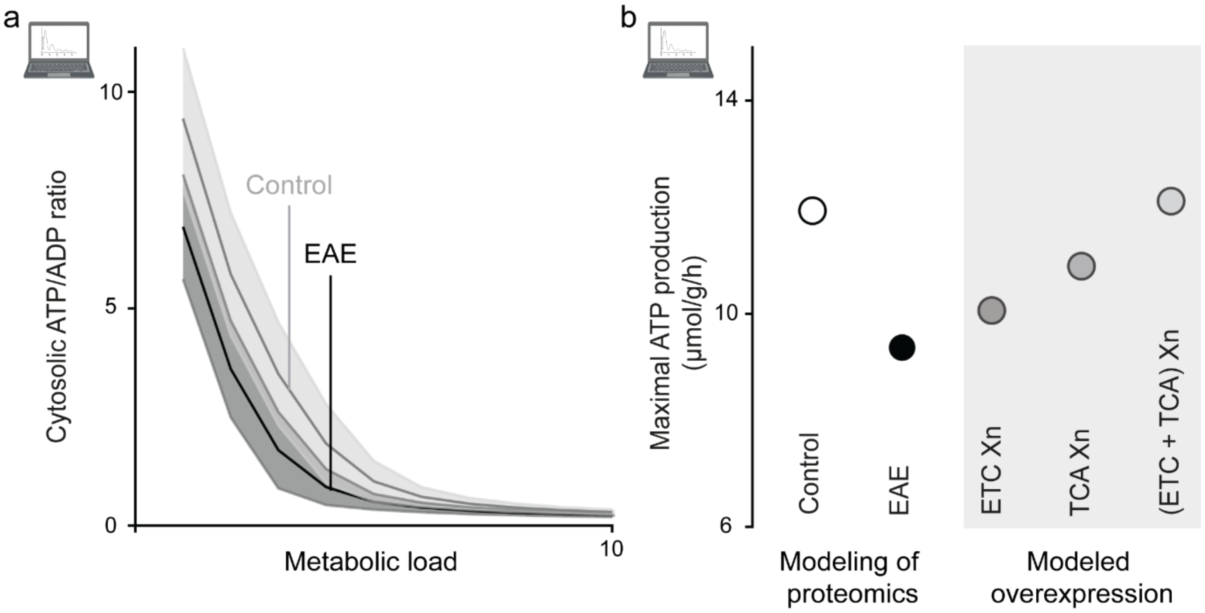


**Tai et al., Extended Data Figure 8**

**Extended Data Figure 8: Prediction of energy state using QSM™-based metabolic profiling.**

(a) Comparisons of predicted maximal ATP production based on AAV/*MitoTag*-based proteomic analysis in control, EAE, modelled overexpression (Xn) of ETC, TCA cycle and both ETC and TCA cycle. (b) Predicted cytosolic ATP/ADP ratio with increasing metabolic load in EAE (dark grey) compared to healthy control (light grey).
